# 3D Structural Connectivity in Healthy Adults: Detecting Structural Connections and Their Underlying Substrates for Olfactory Saccadic Pathway

**DOI:** 10.1101/483628

**Authors:** Ganesh Elumalai, Imran Shareef Syed, Harshita Chatterjee, Valencia Brown, Lekesha Adele Sober, Nitya Akarsha Surya, Venkata Ghanta, Pradeep Chandrasekhar, Anbarasan Venkatesan, Nadira Sewram, Nitesh Raj Muthukumaran, Golla Harkrishna, Nanduri Mojess Vamsi, Nneoma Somtochukwu Osakwe

**Affiliations:** Team NeurON, Texila American University

## Abstract

This confirmatory study aimed to unravel the neural structural connectivity of Olfactory-Saccadic pathways extending between Piriform and Entorhinal Cortices to Frontal Eye Field (FEF), and to correlate its functional importance with possible clinical implications, using Diffusion Imaging fiber Tractography. The confirmatory observational analysis used thirty-two healthy adults, ultra-high b-value, diffusion imaging datasets from an Open access platform in Human Connectome Project (HCP). In all the datasets from both the sexes, fibers were traced and the neural structural connectivity was confirmed. The hemispheric differences between male and female subjects were anaNumber of tracts refers to a bundle of fibres having lysed using independent sample t-test. Thus, the study confirmed the structural existences of Olfactory-saccadic pathways that may be involved in influencing the movements of the neck and eyeball gaze (saccadic eye movement), towards the spatial orientation of olfactory stimulus.

## INTRODUCTION

**S**accadic eye movement can be described as a visual process that can be achieved by orienting the eyes towards an object due to influence from a particular stimulus, a process controlled by CNS ^[1]^. “The olfactory-visual saccadic pathway is a new structural finding observed by our “Team NeurON”. On an attempt to trace the neural structural connectivity of Olfactory-motor pathways, we identified some new structural connections between the “Piriform and Entorhinal Cortex (Brodmann’s area 27, 28 and 34) to Frontal Eye Field (FEF)” (Brodmann’s area 8). The previous evidences on these hypothetical streams remain unclear, hence we pursued these connections to confirm its structural existence, and to identify its possible functional and clinical correlations. Mitral cells or tufted cells in the olfactory bulb receives axons from the olfactory nerve (CN I) and the axons of mitral cells project to the ipsilateral olfactory cortex through the lateral olfactory tract. These Axons project to primary olfactory areas like the anterior olfactory nucleus, piriform cortex, anterior cortical amygdaloid nucleus, peri-amygdaloid cortex and entorhinal cortex ^[2]^. The piriform cortex would integrate the olfactory information from the olfactory bulb and transmit to the prefrontal cortex where FEF is located ^[3]^. The FEF and area 8A is a functional field in the prefrontal cortex, located in the rostral bank of arcuate sulcus as seen in the macaque’s monkey ^[4]^.

FEF participates in the transformation of visual signals into saccades in conjunction with the supplementary eye fields, insula, and median part of the cingulate gyrus ^[4]^. Both the superior colliculus and the frontal eye field (Brodmann’s area 8) are the gaze centres, important for the initiation and accurate targeting of saccadic eye movements ^[5]^. A test initially demonstrated on the monkey shows that there are two types of saccades which activate the FEF in total darkness that gives an idea that all fully volitional saccades are preceded by FEF activation ^[6]^. FEF contains two groups of different neurons; he first group consists of movement neurons that do not respond to visual stimulus directly but are active before and during saccades. They respond in the opposite way to fixation neurons. The second group are visually and aurally responsive neurons that are active during target discrimination independent of saccade programming ^[7]^. According to some references from previous findings, it is mentioned that vision drives olfactory perception, but there has been little indication that olfaction could modulate visual perception ^[8]^. Shenbing Kuang & Tao Zhang performed a test to extend their understandings on vision-olfaction couplings and suggests that a functional interaction between the visual dorsal pathway and the olfactory system exist ^[9]^.

When an odour approaches from an unknown distance and from unknown direction, it can still be perceived without looking hence one can perceive, detect, discriminate and identify a given olfactory stimulus, whether it is coffee, banana or some delicious food etc., then our eyes engage in rapid movement (saccadic eye movement) towards the localized stimulus for fixation. So, according to our hypothesis, we are suggesting that there exists some structural connectivity between the primary olfactory cortex to the frontal eye field and in case of any damage to this connectivity, it may lead to olfactory attention deficit. The olfactory impairments, described in various neurodegenerative and psychiatric disorders involve deficits in the detection, discrimination, and identification of odours, and so they are likely to share affected brain anatomical substrates. Olfactory dysfunction is mainly associated with conditions such as Alzheimer’s disease, Parkinson’s disease (including other diseases caused by Lewy bodies), Autism and Obsessive-ompulsive disorder (OCD), mesial temporal lobe epilepsy, schizophrenia, Huntington’s Disease and multiple sclerosis among others ^[2]^ ^[10]^.

Where, FEF= Frontal Eye Field and PEC= Piriform and Entorhinal Cortex

## RESULTS

Fibre tracking datasets were collected from 32 subjects, data which confirmed the existence of a structural connectivity between the Piriform and Entorhinal cortices and the frontal eye field. The datasets were analysed within the following parameters: (A)-Number of tracts, (B)-Tract volume (mm3), (C)-Tract length mean (mm) and (D)-Tract length standard deviation (mm) and variances within the parameters observed were also evaluated among male and female subjects. Subsiding our principal hypothesis, we theorized that similarities would be observed in the connectivity that exist within the hemispheres in with respect to each sex and also when compared between both sexes.

### A) Number of tracts

Number of tracts refers to a bundle of fibres having a common origin, termination and functions to integrate and generate output. Upon analysis of our supplementary thesis suggesting differences in the connectivity between the hemispheres in male and female subjects, we compared the number of tracts within the following specifications: i) male unilateral and bilateral hemispheric variances, ii) female unilateral and bilateral hemispheric variances and iii) male and female unilateral variances on right side and iv) male and female unilateral variances on left side.

#### i) A. Male unilateral hemispheric variance

Upon analysis of the 16 right side datasets for male subjects (Table-1), we found that the subject MGH_1009 (Fig-1) with the mean age of 32 years have the highest number of tracts (904), and the dataset MGH_1028 (Fig-2) with the mean age of 37 with the least number of tracts (2). Similar observation of the 16 male datasets from the left side (Table-2) showed that the subject MGH_1014 (Fig-3) with the mean age of 27 years have the highest number of tracts (274), and the dataset MGH_1015 (Fig-4) with the mean age of 32 with the least number of tracts (8).

**Table 1:**
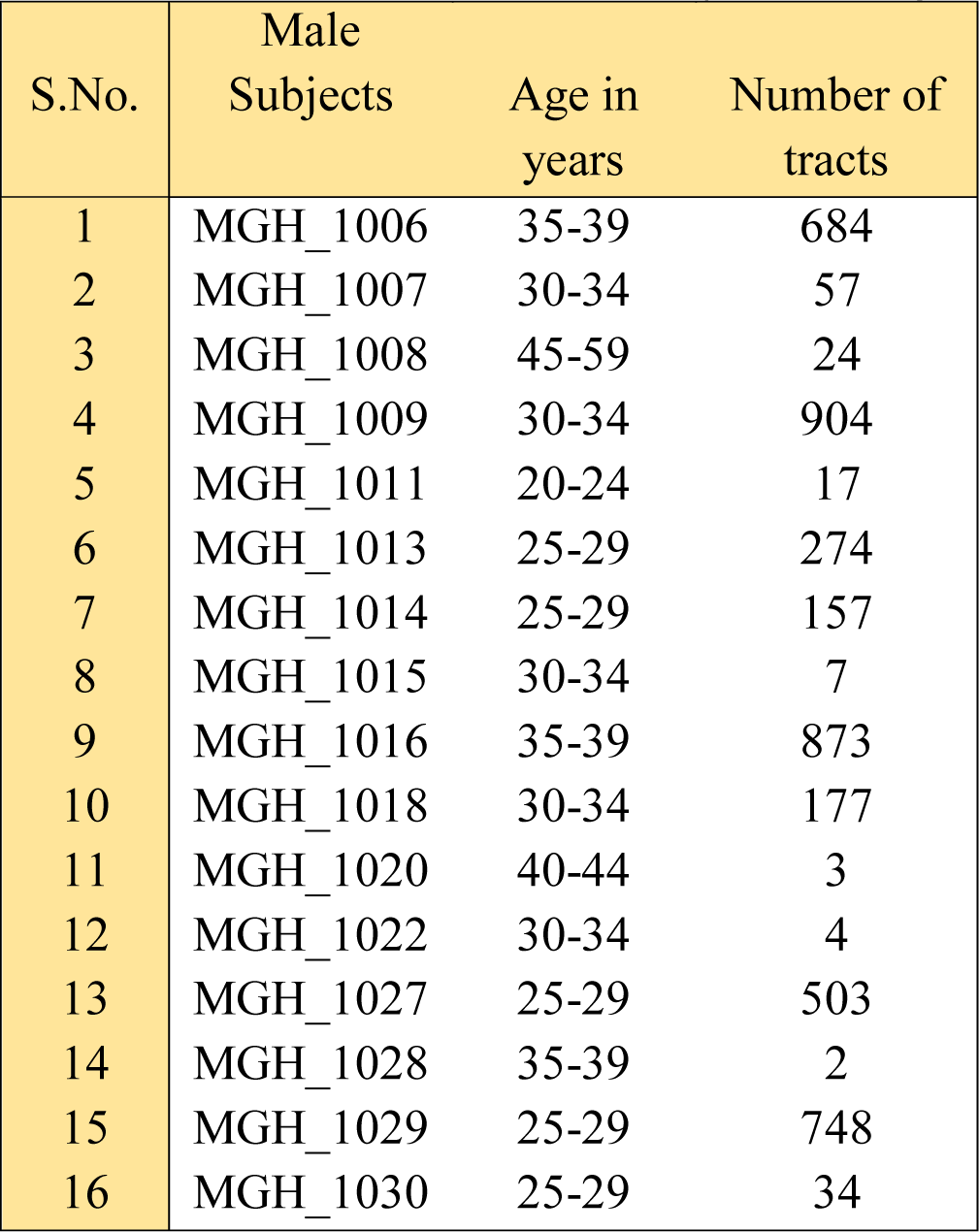
Number of tracts in male subjects on right side

| S.No. | Male Subjects | Age in years | Number of tracts |
| --- | --- | --- | --- |
| 1 | MGH_1006 | 35-39 | 684 |
| 2 | MGH_1007 | 30-34 | 57 |
| 3 | MGH_1008 | 45-59 | 24 |
| 4 | MGH_1009 | 30-34 | 904 |
| 5 | MGH_1011 | 20-24 | 17 |
| 6 | MGH_1013 | 25-29 | 274 |
| 7 | MGH_1014 | 25-29 | 157 |
| 8 | MGH_1015 | 30-34 | 7 |
| 9 | MGH_1016 | 35-39 | 873 |
| 10 | MGH_1018 | 30-34 | 177 |
| 11 | MGH_1020 | 40-44 | 3 |
| 12 | MGH_1022 | 30-34 | 4 |
| 13 | MGH_1027 | 25-29 | 503 |
| 14 | MGH_1028 | 35-39 | 2 |
| 15 | MGH_1029 | 25-29 | 748 |
| 16 | MGH_1030 | 25-29 | 34 |

**Table 2:**
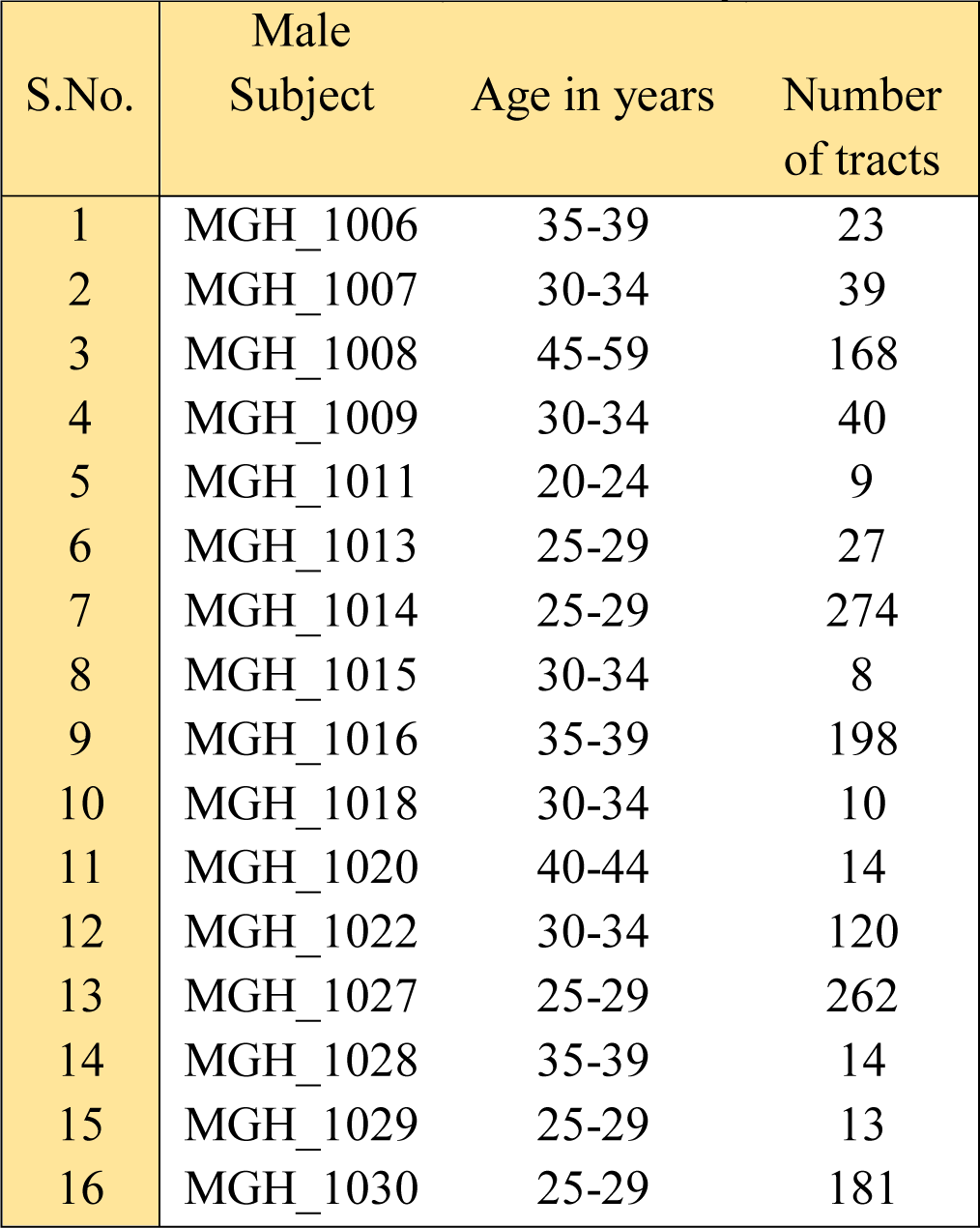
Number of tracts in male subjects on left side

**Fig 1:**
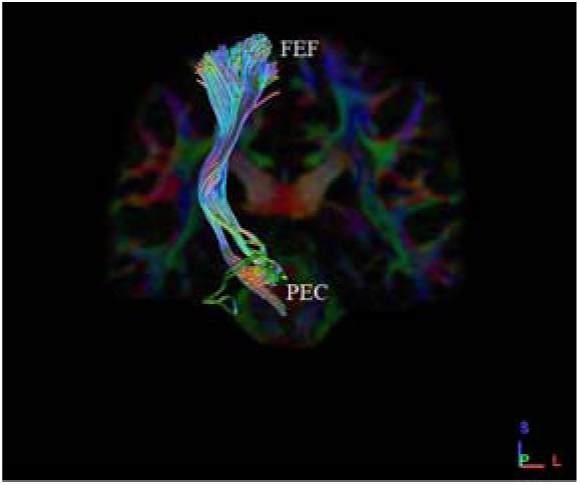
Shows the coronal section with highest (904) number of fibres in male subject (MGH_1009) on right side.

**Fig 2:**
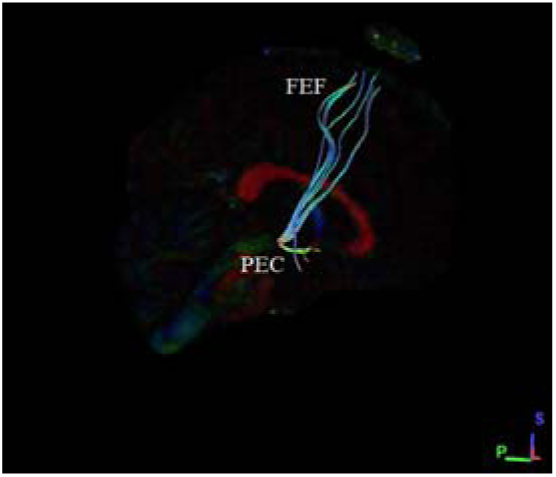
Shows the sagittal section with least number (2) of fibres in male subject (MGH_10028) on right side.

**Fig 3:**
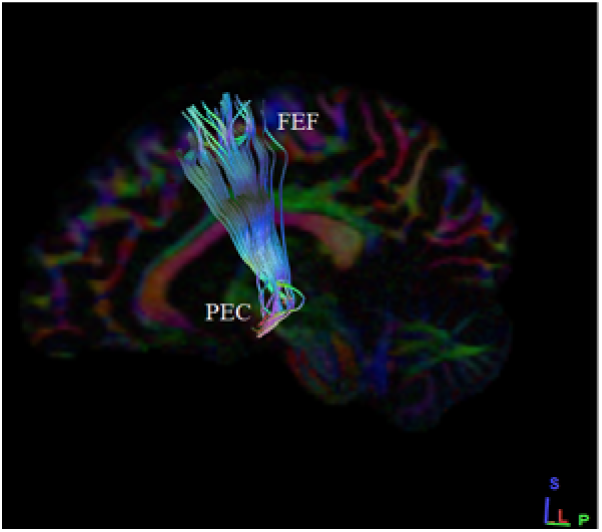
Shows the sagittal section with highest number (274) of fibres in male subject (MGH_1014) on left side.

**Fig 4:**
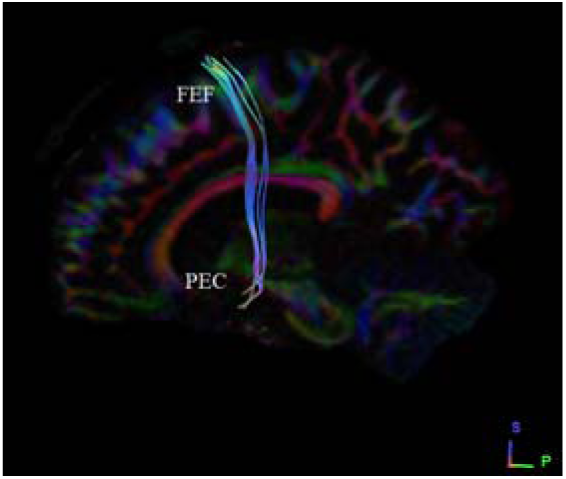
Shows the sagittal section with least number (8) of fibres in male subject (MGH_10015) on left side.

#### i) B. Male bilateral hemispheric variance

An overview observation on bilateral hemispheric variance in male show a greater number of tracts observed in MGH_1009 on the right side (904), but the left side appreciated with only 40 tracts. Similarly, a least number of tracts i.e. 2 are seen on right side in male subject MGH_1028 and 14 on the left side (Table-3).

Upon comparative analysis of male bilateral hemispheric variances, we noted a statistically significant difference among the number of tracts ending in the two hemispheres (Table-3), where fibers exhibited amplified lateralization to the right hemisphere in males (Graph-1), showing deviation from our subsidiary hypothesis which predicted similarities in connectivity between the hemispheres.

**Table 3:**
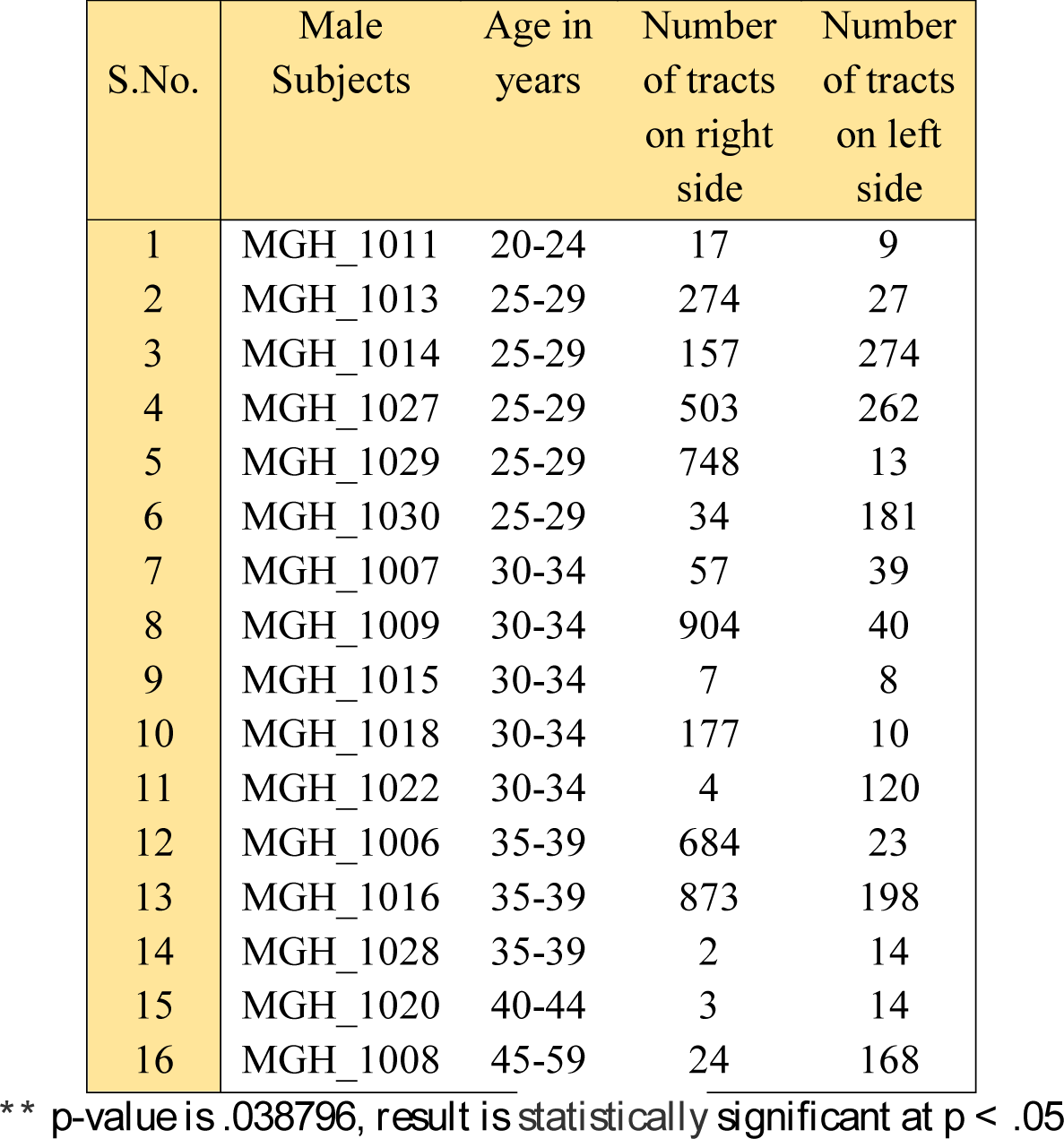
Comparison of number of tracts on male bilateral variances **

| S.No. | Male Subjects | Age in years | Number of tracts on right side | Number of tracts on left side |
| --- | --- | --- | --- | --- |
| 1 | MGH_1011 | 20-24 | 17 | 9 |
| 2 | MGH_1013 | 25-29 | 274 | 27 |
| 3 | MGH_1014 | 25-29 | 157 | 274 |
| 4 | MGH_1027 | 25-29 | 503 | 262 |
| 5 | MGH_1029 | 25-29 | 748 | 13 |
| 6 | MGH_1030 | 25-29 | 34 | 181 |
| 7 | MGH_1007 | 30-34 | 57 | 39 |
| 8 | MGH_1009 | 30-34 | 904 | 40 |
| 9 | MGH_1015 | 30-34 | 7 | 8 |
| 10 | MGH_1018 | 30-34 | 177 | 10 |
| 11 | MGH_1022 | 30-34 | 4 | 120 |
| 12 | MGH_1006 | 35-39 | 684 | 23 |
| 13 | MGH_1016 | 35-39 | 873 | 198 |
| 14 | MGH_1028 | 35-39 | 2 | 14 |
| 15 | MGH_1020 | 40-44 | 3 | 14 |
| 16 | MGH_1008 | 45-59 | 24 | 168 |

\*\* $p$ -value is .038796, result is statistically significant at $p < .05$

**Graph-1:**
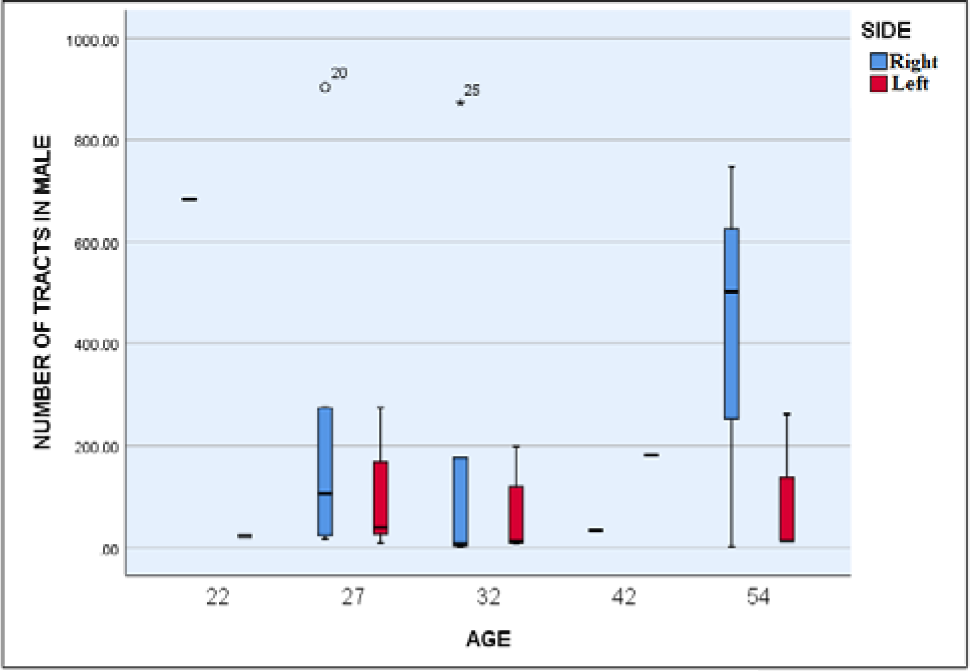
Graphical representation of number of tracts of male bilateral variances.

#### ii) A. Female unilateral hemispheric variance

Analysis of right side data from the 16 female datasets (Table-4) shows that the subject MGH_1033 (Fig-5) with the mean age of 27 years have the highest number of tracts (3025), and the dataset MGH_1019 (Fig-6) with the mean age of 37 presents with the least number of tracts (1).

**Table 4:** Number of tracts in female subjects on right side

| S.No. | Female Subject | Age | Number of tracts |
| --- | --- | --- | --- |
| 1 | MGH_1001 | 35-39 | 288 |
| 2 | MGH_1002 | 30-34 | 204 |
| 3 | MGH_1003 | 45-59 | 1486 |
| 4 | MGH_1004 | 30-34 | 132 |
| 5 | MGH_1005 | 20-24 | 8 |
| 6 | MGH_1010 | 25-29 | 942 |
| 7 | MGH_1012 | 25-29 | 335 |
| 8 | MGH_1017 | 30-34 | 383 |
| 9 | MGH_1019 | 35-39 | 1 |
| 10 | MGH_1021 | 30-34 | 3 |
| 11 | MGH_1023 | 40-44 | 422 |
| 12 | MGH_1024 | 30-34 | 340 |
| 13 | MGH_1026 | 25-29 | 1482 |
| 14 | MGH_1032 | 35-39 | 3 |
| 15 | MGH_1033 | 25-29 | 3025 |
| 16 | MGH_1035 | 25-29 | 715 |

**Fig 5:**
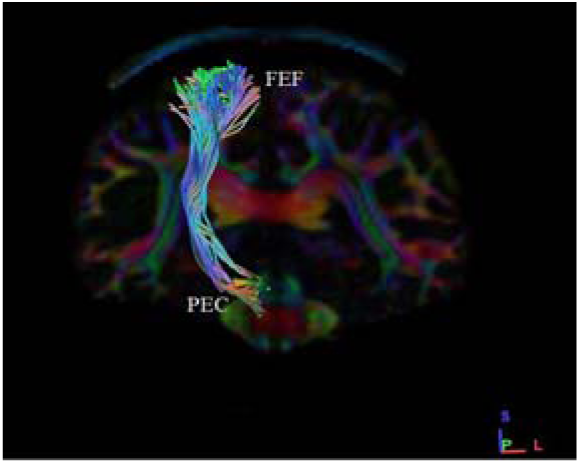
Shows the coronal section with highest number (3025) of fibres in female subject (MGH_1033) on right side.

**Fig 6:**
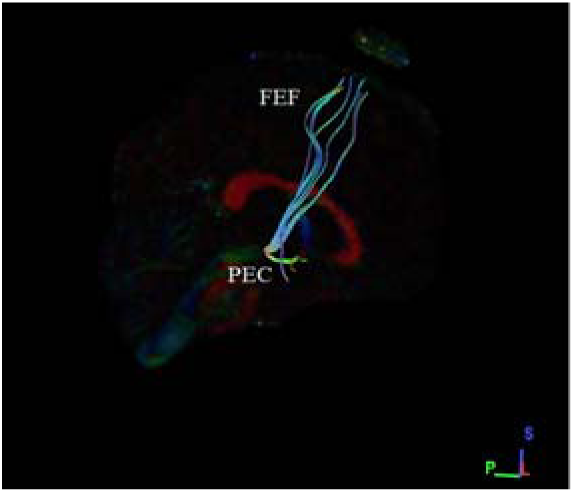
Shows the sagittal section with least number (1) of fibres in female subject (MGH_10019) right on side.

Upon observation of the left side among the 16 female datasets (Table-5), the subject MGH_1026 (Fig-7) with mean age of 27 years have the highest number of tracts (1978), and the dataset MGH_1019 (Fig-8) with the mean age of 37 presents with least number of tracts (6).

**Table 5:** Number of tracts in female subjects on left side

| S.No. | Female Subject | Age | Number of tracts |
| --- | --- | --- | --- |
| 1 | MGH_1001 | 35-39 | 289 |
| 2 | MGH_1002 | 30-34 | 1689 |
| 3 | MGH_1003 | 45-59 | 1490 |
| 4 | MGH_1004 | 30-34 | 84 |
| 5 | MGH_1005 | 20-24 | 7 |
| 6 | MGH_1010 | 25-29 | 129 |
| 7 | MGH_1012 | 25-29 | 715 |
| 8 | MGH_1017 | 30-34 | 28 |
| 9 | MGH_1019 | 35-39 | 6 |
| 10 | MGH_1021 | 30-34 | 18 |
| 11 | MGH_1023 | 40-44 | 408 |
| 12 | MGH_1024 | 30-34 | 125 |
| 13 | MGH_1026 | 25-29 | 1978 |
| 14 | MGH_1032 | 35-39 | 8 |
| 15 | MGH_1033 | 25-29 | 471 |
| 16 | MGH_1035 | 25-29 | 256 |

**Fig 7:**
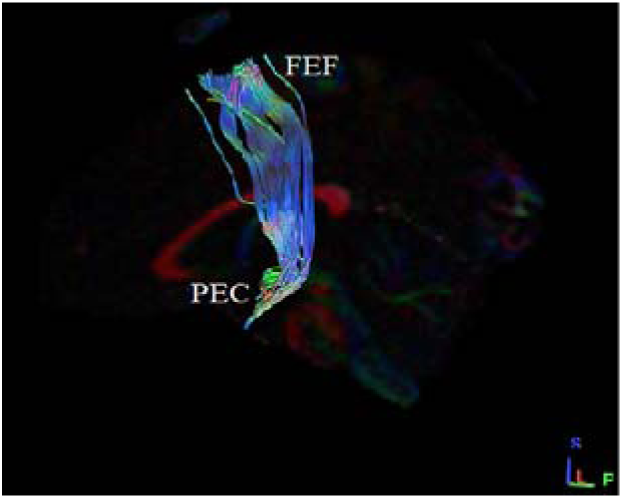
Shows the sagittal section with highest number (1978) of fibres in female subject (MGH_1026) on left side.

**Fig 8:**
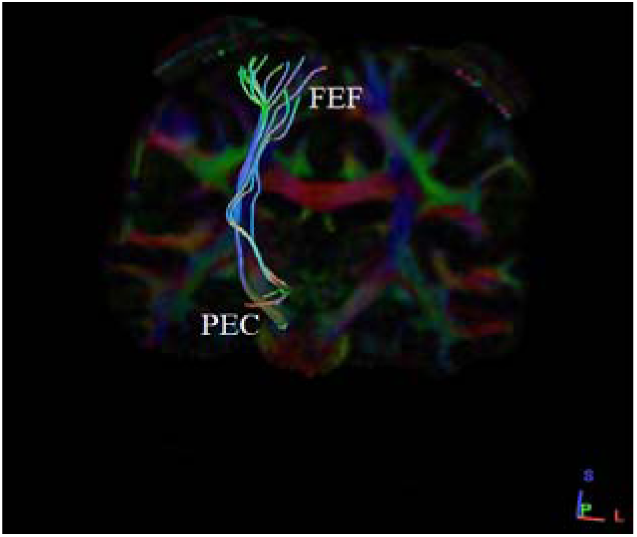
Shows the coronal section with least number (6) of fibres in female subject (MGH_10019) right on left side.

#### ii) B. Female bilateral hemispheric variance

Observation of bilateral hemispheric variances in female shows a greater number of tracts present in MGH_1033 on the right side (3025) with the mean age of 27 years, but the left is with only 471 tracts. Similarly, a least number of tracts i.e. 1 is seen on right side in subject MGH_1019 and 6 on the left side (Table-6).

**Table 6:** Comparison of number of tracts on female bilateral variances *

| S.No. | Female Subject | Age | Number of tracts on right side | Number of tracts on left side |
| --- | --- | --- | --- | --- |
| 1 | MGH_1005 | 20-24 | 8 | 7 |
| 2 | MGH_1010 | 25-29 | 942 | 129 |
| 3 | MGH_1012 | 25-29 | 335 | 715 |
| 4 | MGH_1026 | 25-29 | 1482 | 1978 |
| 5 | MGH_1033 | 25-29 | 3025 | 471 |
| 6 | MGH_1035 | 25-29 | 715 | 256 |
| 7 | MGH_1002 | 30-34 | 204 | 1689 |
| 8 | MGH_1004 | 30-34 | 132 | 84 |
| 9 | MGH_1017 | 30-34 | 383 | 28 |
| 10 | MGH_1021 | 30-34 | 3 | 18 |
| 11 | MGH_1024 | 30-34 | 340 | 125 |
| 12 | MGH_1001 | 35-39 | 288 | 289 |
| 13 | MGH_1019 | 35-39 | 1 | 6 |
| 14 | MGH_1032 | 35-39 | 3 | 8 |
| 15 | MGH_1023 | 40-44 | 422 | 408 |
| 16 | MGH_1003 | 45-59 | 1486 | 1490 |

\*The $p$ -value is .620879, result is statistically not significant at $p < .05$ .

Comparative evaluation of the set of results obtained for female bilateral hemispheric variance showed no significant difference in the number of tracts in both hemispheres (Table-6), similar results were observed for both right and left hemispheres (Graph-2), hence in support of our subsidiary hypothesis.

**Graph-2:**
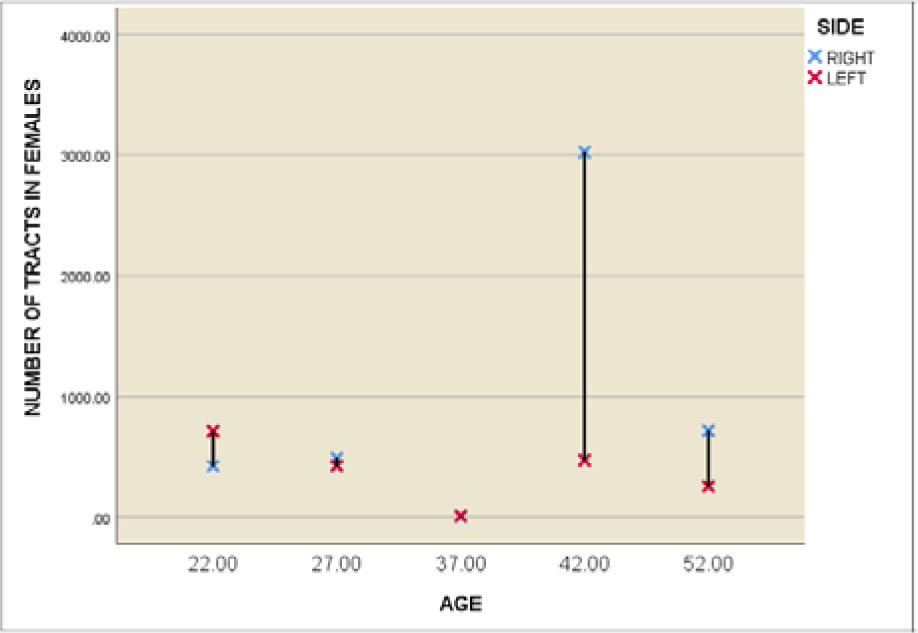
Graphical representation of number of tracts between left and right sides in female subjects.

#### iii) Male and female unilateral hemispheric variances (right side)

Analysis of the 16 right side male datasets (Table-1) showed that the subject MGH_1009 (Fig-1) with the mean age of 32 years have the highest number of tracts (904), and the dataset MGH_1028 (Fig-2) with the mean age of 37 presents with the least number of tracts (2).

Similarly, analysis of the right side for the 16 female datasets (Table-4) showed that the subject MGH_1033 (Fig-5) with the mean age of 27 years have highest number of tracts (3025), and the dataset MGH_1019 (Fig-6) with the mean age of 37 presents with least number of tracts (1).

Correlative observation on unilateral hemispheric variances in male and female subjects on right side is shown in Table-7.

**Table 7:**
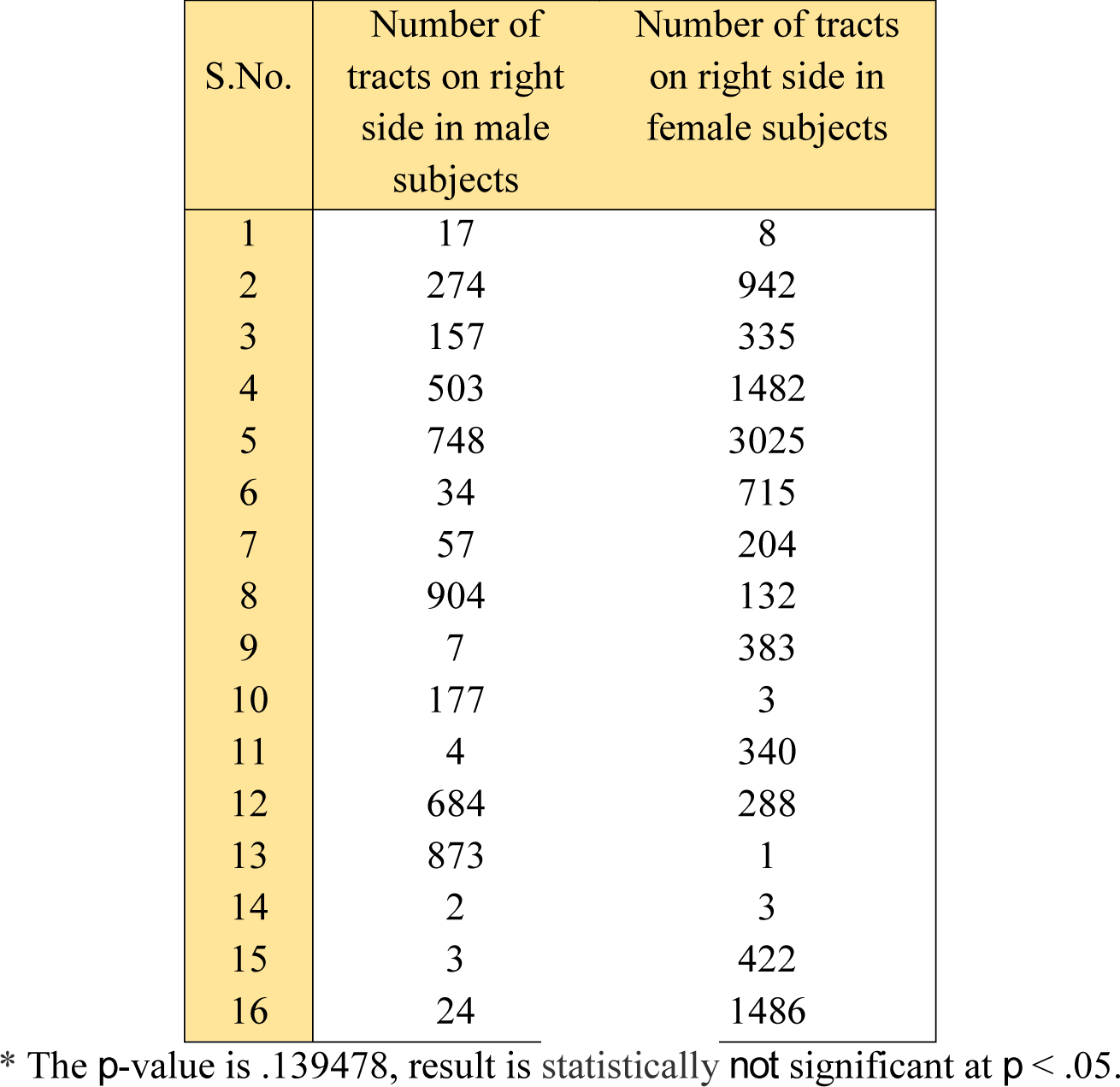
Comparison between number of tracts of both male and female unilateral hemispheric variances on right side *

Conclusively, from our set of results for male and female unilateral hemispheric variances on right side, the results deemed not significant since no substantial difference was seen in the number of tracts in the hemispheres (Table-7) as similar results were observed in both male and female unilateral hemispheres (Graph-3), hence in support of our subsidiary hypothesis.

#### iv) Male and female unilateral hemispheric variances (left side)

Upon observation of the 16 male left side datasets (Table-2), it was seen that the subject MGH_1014 (Fig-3) with the mean age of 27 years have the highest number of tracts (274), and the datasets MGH_1015 (Fig-4) with the mean age of 32 having the least number of tracts (8).

Observation of the left side datasets for the 16 females (Table-5) showed that the subject MGH_1026 (Fig-7) with the mean age of 27 years have highest number of tracts (1978), and then the datasets MGH_1019 (Fig-8) with the mean age of 37 presents with the least number of tracts (6).

Correlative observation on unilateral hemispheric variances of male and female subjects on left side is illustrated in (Table-8).

**Table 8:** Comparison between number of tracts of both male and female unilateral hemispheric

| S.No. | Number of tracts on left side in male subjects | Number of tracts on left side in female subjects |
| --- | --- | --- |
| 1 | 9 | 7 |
| 2 | 27 | 129 |
| 3 | 274 | 715 |
| 4 | 262 | 1978 |
| 5 | 13 | 471 |
| 6 | 181 | 256 |
| 7 | 39 | 1689 |
| 8 | 40 | 84 |
| 9 | 8 | 28 |
| 10 | 10 | 18 |
| 11 | 120 | 125 |
| 12 | 23 | 289 |
| 13 | 198 | 6 |
| 14 | 14 | 8 |
| 15 | 14 | 408 |
| 16 | 168 | 1490 |

\*\* The $p$ -value is .023261, result is statistically significant at $p < .05$ .

Conclusively, from our set of results for male and female unilateral hemispheric variances on left side, we noted that a statistically significant difference exists in the number of tracts in the hemispheres (Table-8), with a demonstration of amplified lateralization to the left hemisphere in females (Graph-4), showing deviation from our subsidiary hypothesis which predicted similarities in connectivity between the hemispheres.

**Graph-3:**
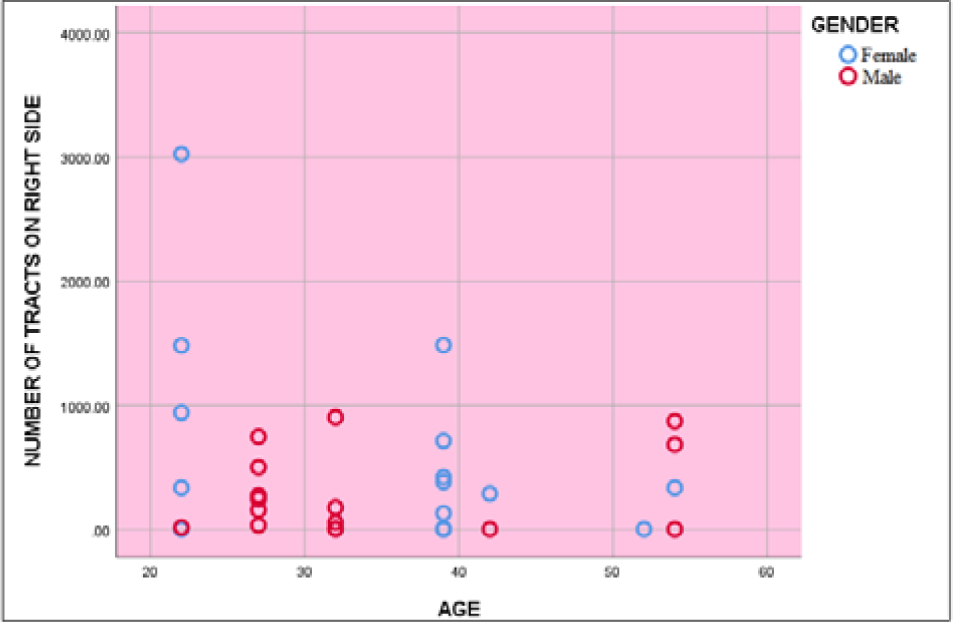
Graphical representation of number of tracts on male and female subjects on right side.

**Graph-4:**
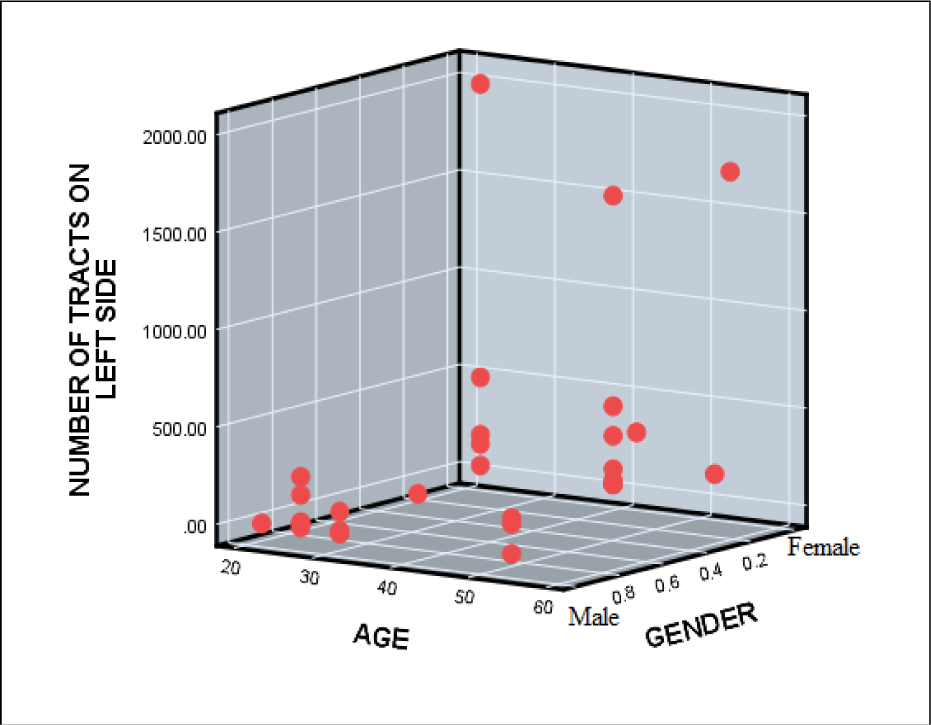
Graphical representation of number of tracts on male and female subjects on left side.

### B) Tract volume (mm^3^)

Tract volume is the total volume occupied by each fibre with the function to integrate and generate neural output. We analyzed the tract volume (mm3) present in both the sexes i.e. i) male subjects and ii) female subjects and compared the variances observed in connectivity between the two hemispheres.

#### i) A. Male unilateral hemispheric variance

Upon analysis of the right side for the 16 male datasets (Table-9), we found that the subject MGH_1009 (Fig-9) with the mean age of 32 years have the highest tract volume (mm3) (15136.9), and the dataset MGH_1028 (Fig-10) with the mean age of 37 presents with least tract volume (mm3) (523.13).

**Table 9:** Tract volume (mm^3^) in male subjects on right side

| S.No. | Male Subject | Age in years | Tract volume (mm<sup>3</sup>) |
| --- | --- | --- | --- |
| 1 | MGH_1006 | 35-39 | 13263.70 |
| 2 | MGH_1007 | 30-34 | 6024.38 |
| 3 | MGH_1008 | 45-59 | 1704.38 |
| 4 | MGH_1009 | 30-34 | 15136.9 |
| 5 | MGH_1011 | 20-24 | 2021.62 |
| 6 | MGH_1013 | 25-29 | 8390.25 |
| 7 | MGH_1014 | 25-29 | 8478.00 |
| 8 | MGH_1015 | 30-34 | 1144.13 |
| 9 | MGH_1016 | 35-39 | 8775.00 |
| 10 | MGH_1018 | 30-34 | 9234.00 |
| 11 | MGH_1020 | 40-44 | 722.25 |
| 12 | MGH_1022 | 30-34 | 1228.50 |
| 13 | MGH_1027 | 25-29 | 11637.00 |
| 14 | MGH_1028 | 35-39 | 523.13 |
| 15 | MGH_1029 | 25-29 | 9952.87 |
| 16 | MGH_1030 | 25-29 | 1957.50 |

**Fig 9:**
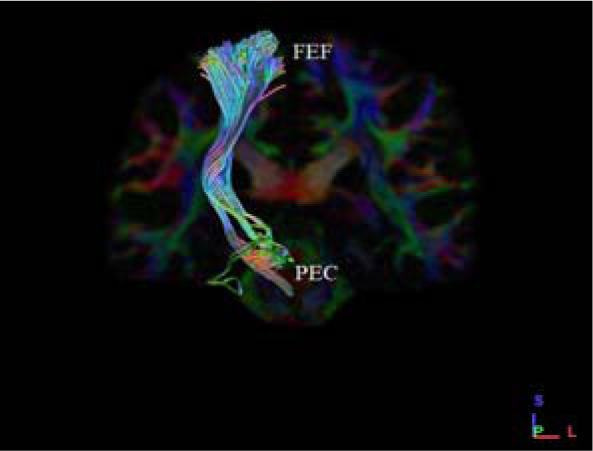
Shows the coronal section with highest (15136.9) tract volume (mm^3^) in male subject (MGH_1009) on right side.

**Fig 10:**
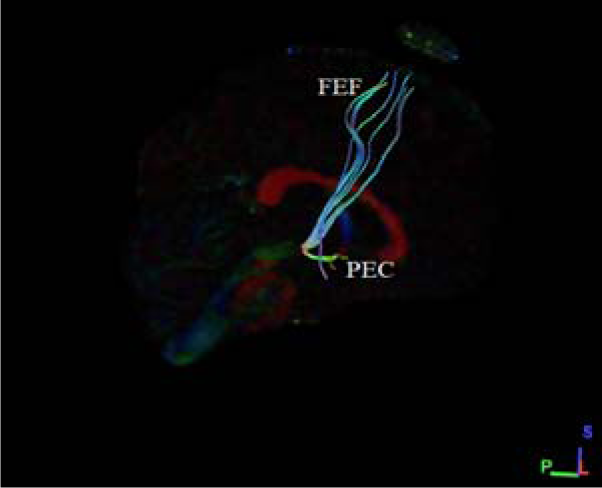
Shows the sagittal section with least (523.13) Tract volume (mm^3^) in male subject (MGH_1028) on right side.

From observations of the left side for the 16 male datasets (Table-10), it was seen that the subject MGH_1014 (Fig-11) with the mean age of 27 years have the highest tract volume (mm3) (9625.50), and the dataset MGH_1015 (Fig-12) with the mean age of 32 presents with least tract volume (mm3) (1275.75).

**Table 10:** Tract volume (mm^3^) in male subjects on left side

| S.No. | Male Subject | Age in years | Tract volume (mm<sup>3</sup>) |
| --- | --- | --- | --- |
| 1 | MGH_1006 | 35-39 | 1873.12 |
| 2 | MGH_1007 | 30-34 | 4448.25 |
| 3 | MGH_1008 | 45-59 | 5217.75 |
| 4 | MGH_1009 | 30-34 | 5504.62 |
| 5 | MGH_1011 | 20-24 | 1373.62 |
| 6 | MGH_1013 | 25-29 | 2521.13 |
| 7 | MGH_1014 | 25-29 | 9625.50 |
| 8 | MGH_1015 | 30-34 | 1275.75 |
| 9 | MGH_1016 | 35-39 | 4249.12 |
| 10 | MGH_1018 | 30-34 | 1285.87 |
| 11 | MGH_1020 | 40-44 | 1292.62 |
| 12 | MGH_1022 | 30-34 | 6284.25 |
| 13 | MGH_1027 | 25-29 | 4357.12 |
| 14 | MGH_1028 | 35-39 | 3796.88 |
| 15 | MGH_1029 | 25-29 | 2669.62 |
| 16 | MGH_1030 | 25-29 | 4158.00 |

**Fig 11:**
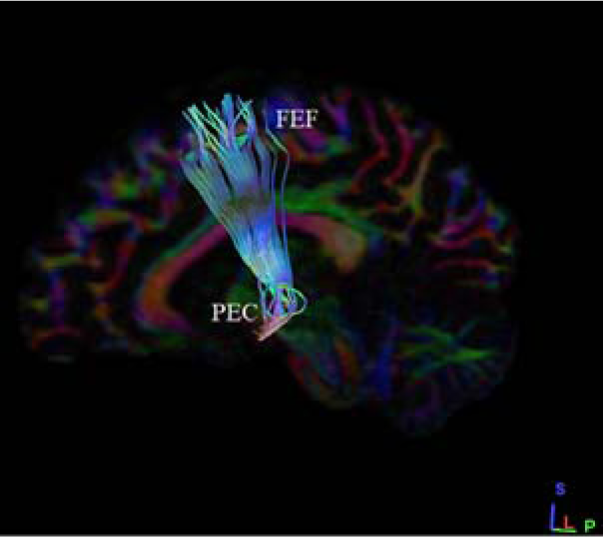
Shows the sagittal section with highest (9625.50) Tract volume (mm^3^) in male subject (MGH_1014) on left side.

**Fig 12:**
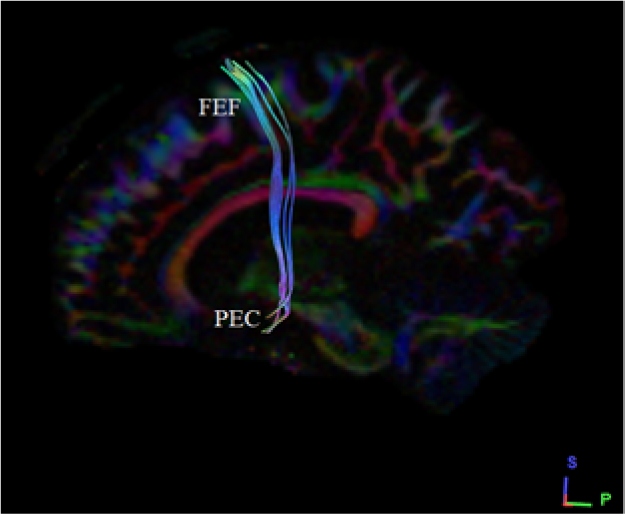
Shows the sagittal section with least (1275.75) Tract volume (mm^3^) in male subject (MGH_10015) on left side.

#### i) B. Male bilateral hemispheric variance

An overview observation of bilateral hemispheric variance in male showed a greater tract volume (mm3) observed in MGH_1009 on the right side (15136.9), but the left appreciated with only 5504.62 tract volume. Similarly, a least tract volume (mm3) i.e. 523.13 is seen on right side in male subject MGH_1028 and 3796.88 on his left side (Table-11).

**Table 11:** Comparison of tract volume (mm^3^) on male bilateral variances *

| S.No. | Male Subjects | Age in years | Tract volume (mm<sup>3</sup>) on right side | Tract volume (mm<sup>3</sup>) on left side |
| --- | --- | --- | --- | --- |
| 1 | MGH_1011 | 20-24 | 2021.62 | 1373.62 |
| 2 | MGH_1013 | 25-29 | 8390.25 | 2521.13 |
| 3 | MGH_1014 | 25-29 | 8478.00 | 9625.50 |
| 4 | MGH_1027 | 25-29 | 11637.00 | 4357.12 |
| 5 | MGH_1029 | 25-29 | 9952.87 | 2669.62 |
| 6 | MGH_1030 | 25-29 | 1957.50 | 4158.00 |
| 7 | MGH_1007 | 30-34 | 6024.38 | 4448.25 |
| 8 | MGH_1009 | 30-34 | 15136.9 | 5504.62 |
| 9 | MGH_1015 | 30-34 | 1144.13 | 1275.75 |
| 10 | MGH_1018 | 30-34 | 9234.00 | 1285.87 |
| 11 | MGH_1022 | 30-34 | 1228.50 | 6284.25 |
| 12 | MGH_1006 | 35-39 | 13263.70 | 1873.12 |
| 13 | MGH_1016 | 35-39 | 8775.00 | 4249.12 |
| 14 | MGH_1028 | 35-39 | 523.13 | 3796.88 |
| 15 | MGH_1020 | 40-44 | 722.25 | 1292.62 |
| 16 | MGH_1008 | 45-59 | 1704.38 | 5217.75 |

\* The $p$ -value is .074467, result is statistically *not* significant at $p < .05$ .

Conclusively, for the set of results observed in male bilateral hemispheric variance, we noted no significant difference exists in the tract volume (mm3) in the two hemispheres (Table-11), similar results were seen from both right and left hemispheres (Graph-5), hence supporting our subsidiary hypothesis.

**Graph-5:**
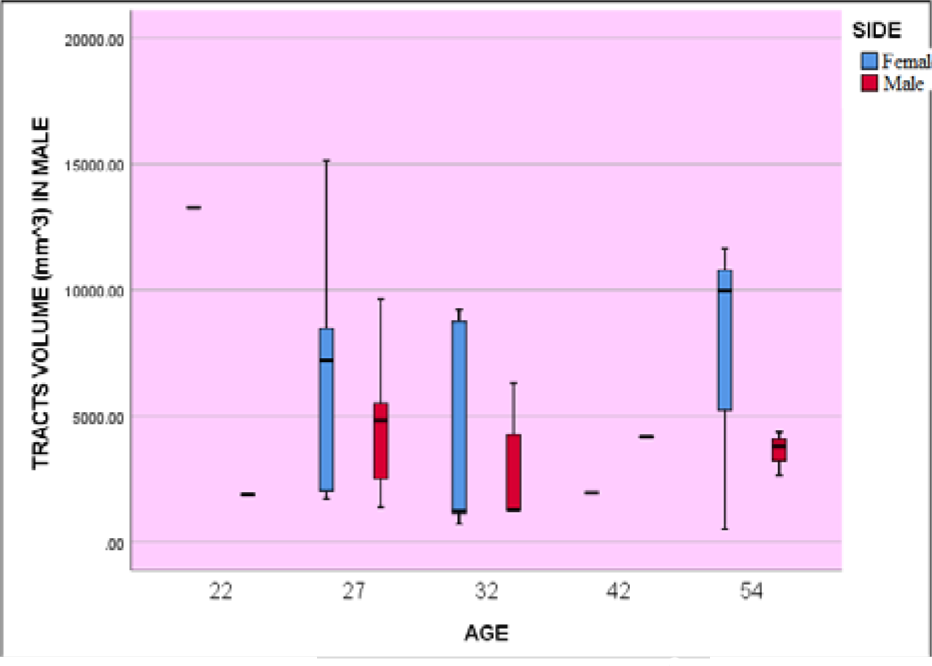
Graphical representation of tract volume (mm^3^) between left and right sides in male subjects.

#### ii) A. Female unilateral hemispheric variance

Similarly, upon analysis of the 16 right side datasets for female (Table-12), we found that the subject MGH_1033 (Fig-13) with the mean age of 27 years have the highest tract volume (mm3) (18245.20), and the dataset MGH_1019 (Fig-14) with the mean age of 37 have least tract volume (mm3) (131.63).

**Table 12:**
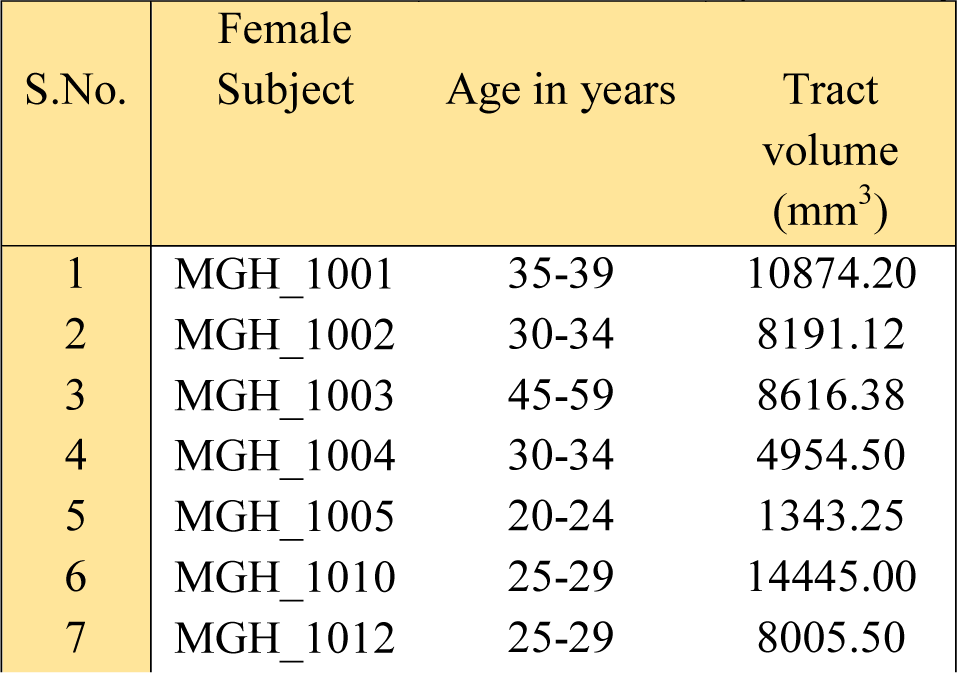
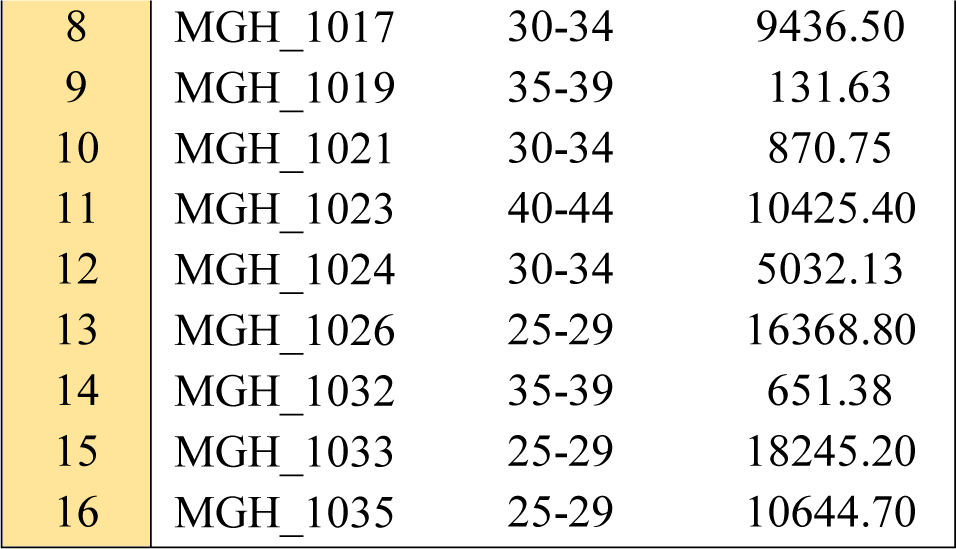
Tract volume (mm^3^) in female subjects on right side

**Fig 13:**
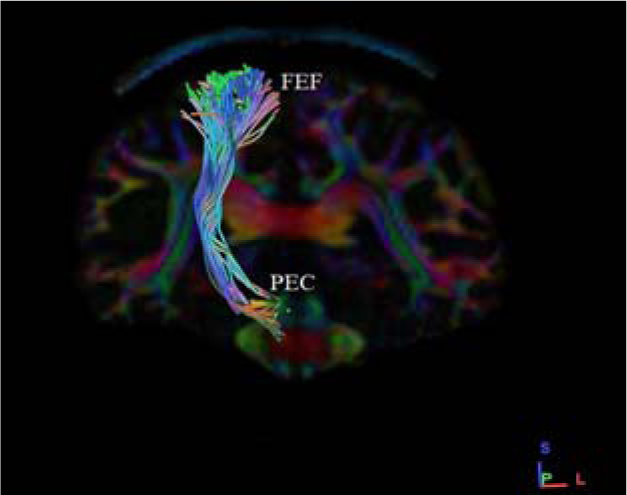
Shows the coronal section with highest (1824.20) Tract volume (mm^3^) in female subject (MGH_1033) on right side.

**Fig 14:**
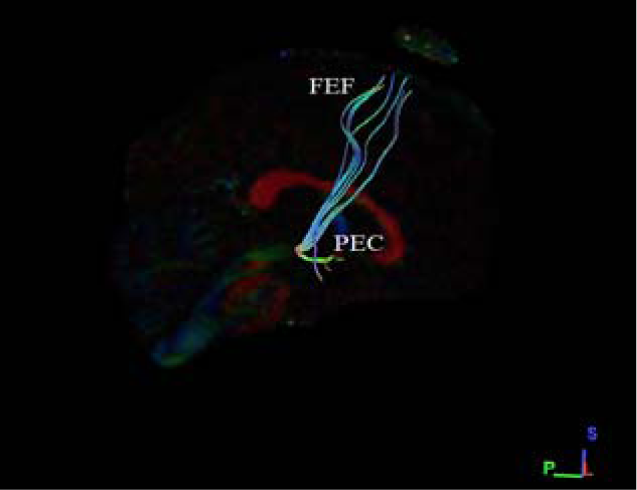
Shows the coronal section with least (131.63) Tract volume (mm^3^) in female subject (MGH_1019) right on right side.

Upon observation of the 16 female left side datasets (Table-13), it was seen that the subject MGH_1026 (Fig-15) with the mean age of 27 years have the highest tract volume (mm3) (13311.00), and the dataset MGH_1019 (Fig-16) with the mean age of 37 have least tract volume (mm3) (1107.00).

**Table 13:** Tract volume (mm^3^) in female subjects on left side

| S.No. | Female Subject | Age in years | Tract volume (mm<sup>3</sup>) |
| --- | --- | --- | --- |
| 1 | MGH_1001 | 35-39 | 7263.00 |
| 2 | MGH_1002 | 30-34 | 9986.62 |
| 3 | MGH_1003 | 45-59 | 8633.25 |
| 4 | MGH_1004 | 30-34 | 2149.87 |
| 5 | MGH_1005 | 20-24 | 1420.87 |
| 6 | MGH_1010 | 25-29 | 9429.75 |
| 7 | MGH_1012 | 25-29 | 5791.50 |
| 8 | MGH_1017 | 30-34 | 2143.12 |
| 9 | MGH_1019 | 35-39 | 1107.00 |
| 10 | MGH_1021 | 30-34 | 2922.75 |
| 11 | MGH_1023 | 40-44 | 7543.13 |
| 12 | MGH_1024 | 30-34 | 4914.00 |
| 13 | MGH_1026 | 25-29 | 13311.00 |
| 14 | MGH_1032 | 35-39 | 1339.87 |
| 15 | MGH_1033 | 25-29 | 8184.37 |
| 16 | MGH_1035 | 25-29 | 5815.12 |

**Fig-15:**
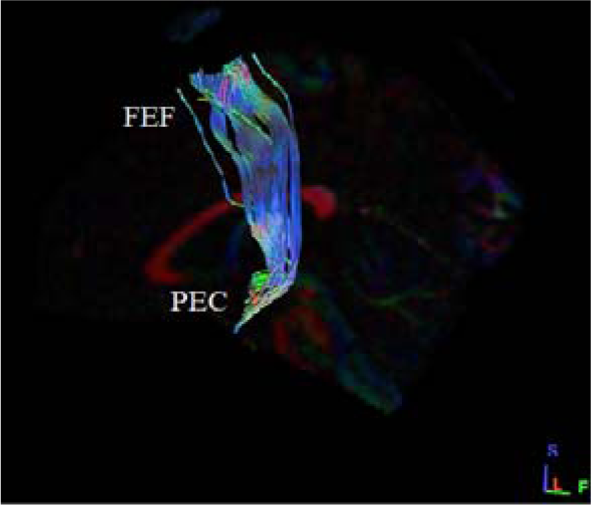
Shows the sagittal section with highest (13311) Tract volume (mm^3^) in female subject (MGH_1026) on left side.

**Fig-16:**
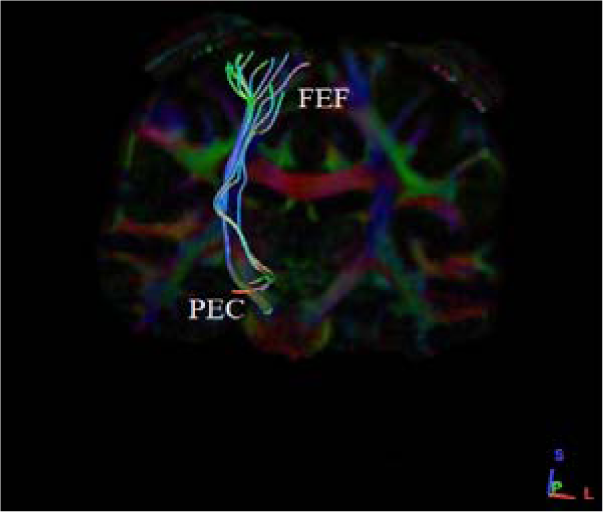
Shows the coronal section with least (1170) Tract volume (mm^3^) in female subject (MGH_10019) right on left side.

#### ii) B. Female bilateral hemispheric variance

An overview observation on bilateral hemispheric variance in female showed a greater tract volume (mm3) observed in MGH_1033 on the right side (18245.20), but the left with only 8184.37 tract volume (mm3). Similarly, a least tract volume (mm3) i.e. 131.63 is seen on right side in female subject MGH_1019 and 1107.00 on the left side (Table-14).

**Table 14:** Comparison of Tract volume (mm^3^) on female bilateral variances *

| S.No. | Female Subject | Age | Tract volume ($\text{mm}^3$) on right side | Tract volume ($\text{mm}^3$) on left side |
| --- | --- | --- | --- | --- |
| 1 | MGH_1005 | 20-24 | 1343.25 | 1420.87 |
| 2 | MGH_1010 | 25-29 | 14445.00 | 9429.75 |
| 3 | MGH_1012 | 25-29 | 8005.50 | 5791.50 |
| 4 | MGH_1026 | 25-29 | 16368.80 | 13311.0 |
| 5 | MGH_1033 | 25-29 | 18245.20 | 8184.37 |
| 6 | MGH_1035 | 25-29 | 10644.70 | 5815.12 |
| 7 | MGH_1002 | 30-34 | 8191.12 | 9986.62 |
| 8 | MGH_1004 | 30-34 | 4954.50 | 2149.87 |
| 9 | MGH_1017 | 30-34 | 9436.50 | 2143.12 |
| 10 | MGH_1021 | 30-34 | 870.75 | 2922.75 |
| 11 | MGH_1024 | 30-34 | 5032.13 | 4914.00 |
| 12 | MGH_1001 | 35-39 | 10874.20 | 7263.00 |
| 13 | MGH_1019 | 35-39 | 131.63 | 1107.00 |
| 14 | MGH_1032 | 35-39 | 651.38 | 1339.87 |
| 15 | MGH_1023 | 40-44 | 10425.40 | 7543.13 |
| 16 | MGH_1003 | 45-59 | 8616.38 | 8633.25 |

\* The $p$ -value is .185525, result is statistically *not* significant at $p < .05$ .

Conclusively from our set of results in female bilateral hemispheric variance, no significant difference exists in the tract volume (mm3) in the two hemispheres (Table-14), similar results were observed in both right and left hemispheres (Graph-6) hence in support of our subsidiary conjecture.

**Graph-6:**
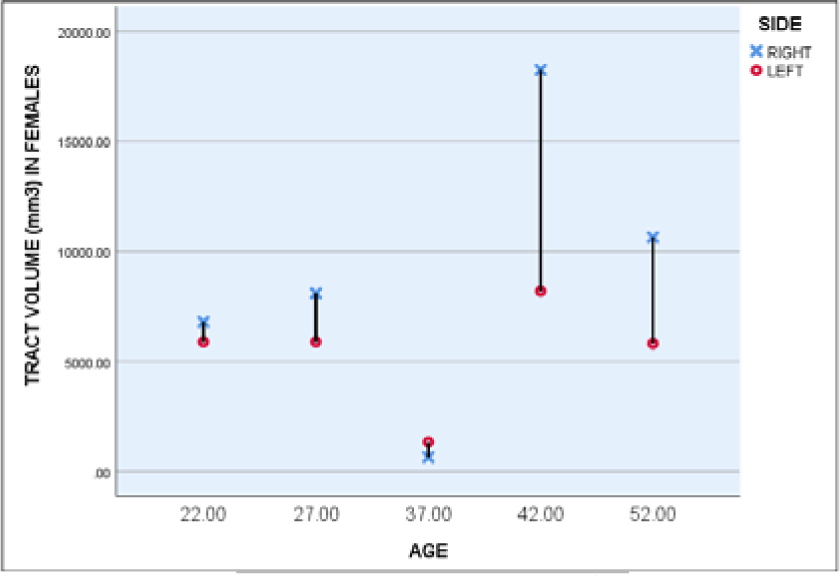
Graphical representation of tract volume (mm^3^) between left and right sides in female subjects.

#### iii) Male and female unilateral hemispheric variances (right side)

Upon analysis of 16 male datasets of right side (Table-9), we found that the subject MGH_1009 (Fig-9) with the mean age of 32 years have highest tract volume (mm3) (15136.9), and the dataset MGH_1028 (Fig-10) with the mean age of 37 having least tract volume (mm3) (523.13).

Similarly, upon analysis of right side for 16 female datasets (Table-12), we found that the subject MGH_1033 (Fig-13) with the mean age of 27 years have the highest tract volume (mm3) (18245.20), and the dataset MGH_1019 (Fig-14) with the mean age of 37 with least tract volume (mm3) (131.63).

Conclusively, for the set of results observed in male and female unilateral hemispheric variances on right side, we noted that no difference exists in the tract volume (mm3) in the hemispheres (Table-15), similar results were observed in both male and female unilateral hemispheres (Graph-7), upholding our subsidiary hypothesis.

**Table 15:** Comparison between Tract volume (mm^3^) of both male and female unilateral hemispheric variances on right side*

| S.No. | Tract volume (mm<sup>3</sup>) on right side in male subjects | Tract volume (mm<sup>3</sup>) on right side in female subjects |
| --- | --- | --- |
| 1 | 2021.62 | 1343.25 |
| 2 | 8390.25 | 14445.00 |
| 3 | 8478.00 | 8005.50 |
| 4 | 11637.00 | 16368.80 |
| 5 | 9952.87 | 18245.20 |
| 6 | 1957.50 | 10644.70 |
| 7 | 6024.38 | 8191.12 |
| 8 | 15136.9 | 4954.50 |
| 9 | 1144.13 | 9436.50 |
| 10 | 9234.00 | 870.75 |
| 11 | 1228.50 | 5032.13 |
| 12 | 13263.70 | 10874.20 |
| 13 | 8775.00 | 131.63 |
| 14 | 523.13 | 651.38 |
| 15 | 722.25 | 10425.40 |
| 16 | 1704.38 | 8616.38 |

\* The p-value is .355377, result is statistically not significant at $p < .05$ .

Correlative observation on unilateral hemispheric variances in male and female on right side is shown in Table-16.

**Table 16:** Comparison between Tract volume (mm^3^) of both male and female unilateral hemispheric variances on left side*

| S.No. | Tract volume (mm<sup>3</sup>) on left side in male subjects | Tract volume (mm<sup>3</sup>) on left side in female subjects |
| --- | --- | --- |
| 1 | 1373.62 | 1420.87 |
| 2 | 2521.13 | 9429.75 |
| 3 | 9625.50 | 5791.50 |
| 4 | 4357.12 | 13311.00 |
| 5 | 2669.62 | 8184.37 |
| 6 | 4158.00 | 5815.12 |
| 7 | 4448.25 | 9986.62 |
| 8 | 5504.62 | 2149.87 |
| 9 | 1275.75 | 2143.12 |
| 10 | 1285.87 | 2922.75 |
| 11 | 6284.25 | 4914.00 |
| 12 | 1873.12 | 7263.00 |
| 13 | 4249.12 | 1107.00 |
| 14 | 3796.88 | 1339.87 |
| 15 | 1292.62 | 7543.13 |
| 16 | 5217.75 | 8633.25 |

\* The p-value is .073925, result is statistically not significant at $p < .05$ .

#### iv) Male and female unilateral hemispheric variances (left side)

Upon observation of the left side in the 16 male datasets (Table-10), it was seen that the subject MGH_1014 (Fig-11) with the mean age of 27 years have the highest tract volume (mm3) (9625.50), and the dataset MGH_1015 (Fig-12) with the mean age of 32 having the least tract volume (mm3) (1275.75).

Upon observation of the left side in the 16 female datasets (Table-14), the subject MGH_1026 (Fig-15) with the mean age of 27 years had the highest tract volume (mm3) (13311.00), and the dataset MGH_1019 (Fig-16) with the mean age of 37 with least tract volume (mm3) (1107.00).

Conclusively, for the set of results observed in male and female unilateral hemispheric variances on left side, we noted no difference exists in the tract volume (mm3) in the hemispheres (Table-16), similar results were observed in both male and female unilateral hemispheres (Graph-8), hence in support of our subsidiary hypothesis.

Correlative observation on unilateral hemispheric variances in male and female on left side is shown in Table-16.

**Graph-7:**
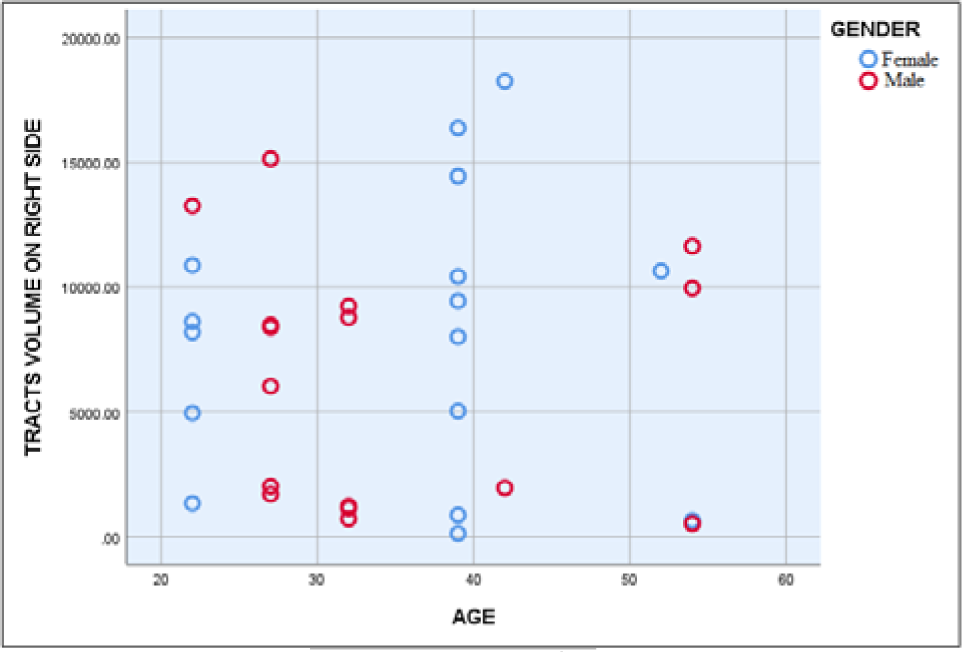
Graphical representation of volume (mm^3^) on male and female subjects on right side.

**Graph-8:**
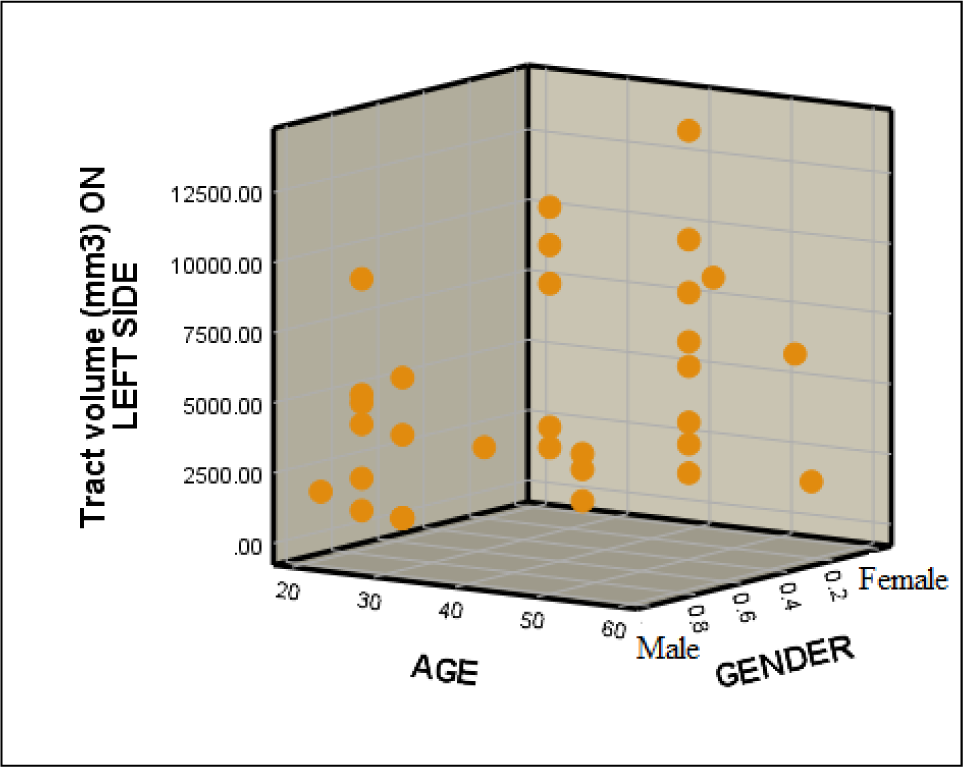
Graphical representation of tract volume (mm^3^) on male and female subjects on left side.

### C) Tract mean length (mm)

Tract mean length (mm), the average absolute deviation of a data set is the average of the absolute deviations from a central point. We analyzed the tract mean length (mm) present in both the sexes i.e. i) male subjects and ii) female subjects.

#### i) A. Male unilateral hemispheric variance

The 16 male datasets analysed from the right side (Table-17) showed that the subject MGH_1022 (Fig-17) with the mean age of 42 years have the highest tract length mean (mm) (152.26), and the dataset MGH_1006 (Fig-18) with the mean age of 37 presents least tract length mean (mm) (85.78).

**Fig 17:**
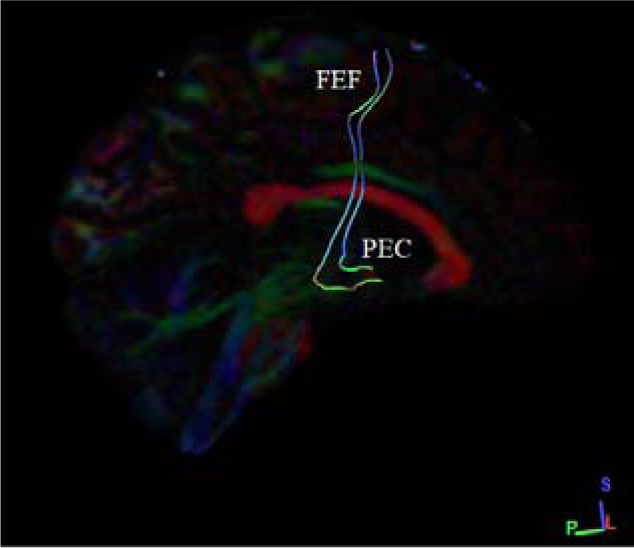
Shows the sagittal section with highest (152.26) Tract mean length (mm) in male subject (MGH_1022) on right side.

**Fig 18:**
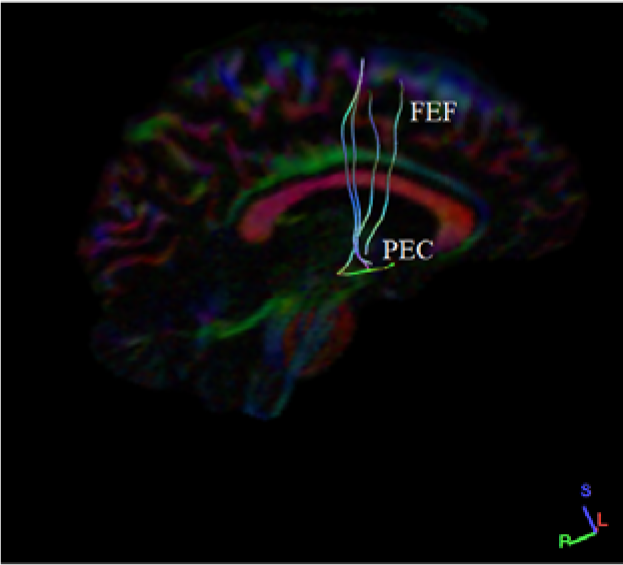
Shows the coronal section with least (85.78) Tract mean length (mm) in male subject (MGH_1006) right on right side.

**Table 17:**
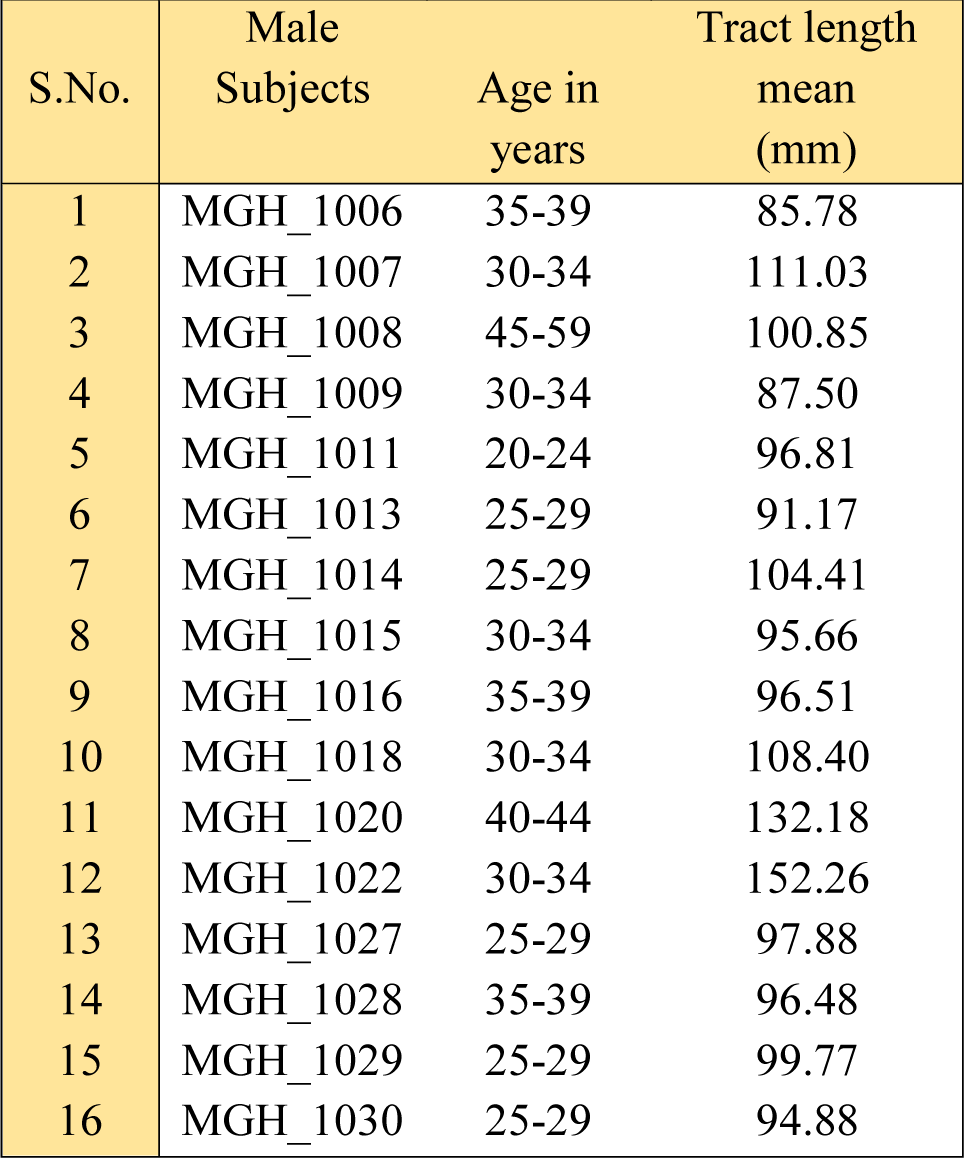
Tract mean length (mm) in male subjects on right side

Upon observation of 16 male datasets from left side (Table-18), it was seen that the subject MGH_1014 (Fig-19) with the mean age of 27 years have the highest tract mean length (mm) (9625.50), and the dataset MGH_1015 (Fig-20) with the mean age of 32 having the least tract mean length (mm) (1275.75).

**Fig 19:**
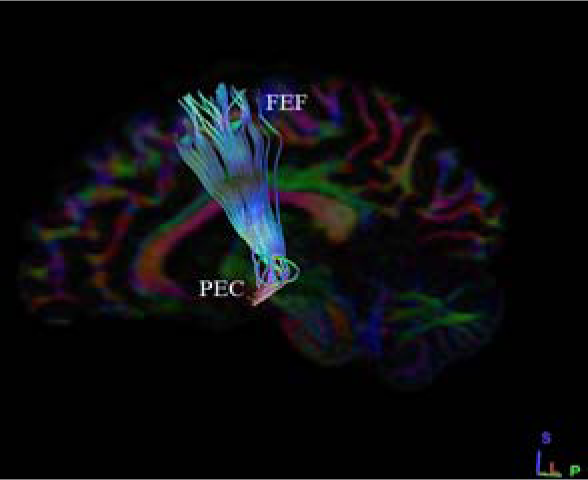
Shows the sagittal section with highest (9625.50) Tract mean length (mm) in male subject (MGH_1014) on left side.

**Fig 20:**
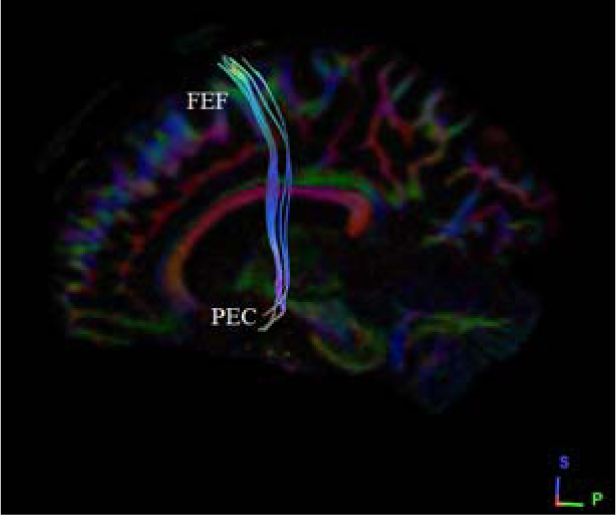
Shows the sagittal section with least (1275.75) Tract mean length (mm) in male subject (MGH_10015) on left side.

**Table 18:** Tract mean length (mm) in male subjects on left side

| S.No. | Male Subjects | Age in years | Tract length mean (mm) |
| --- | --- | --- | --- |
| 1 | MGH_1006 | 35-39 | 1873.12 |
| 2 | MGH_1007 | 30-34 | 4448.25 |
| 3 | MGH_1008 | 45-59 | 5217.75 |
| 4 | MGH_1009 | 30-34 | 5504.62 |
| 5 | MGH_1011 | 20-24 | 1373.62 |
| 6 | MGH_1013 | 25-29 | 2521.13 |
| 7 | MGH_1014 | 25-29 | 9625.50 |
| 8 | MGH_1015 | 30-34 | 1275.75 |
| 9 | MGH_1016 | 35-39 | 4249.12 |
| 10 | MGH_1018 | 30-34 | 1285.87 |
| 11 | MGH_1020 | 40-44 | 1292.62 |
| 12 | MGH_1022 | 30-34 | 6284.25 |
| 13 | MGH_1027 | 25-29 | 4357.12 |
| 14 | MGH_1028 | 35-39 | 3796.88 |
| 15 | MGH_1029 | 25-29 | 2669.62 |
| 16 | MGH_1030 | 25-29 | 4158.00 |

#### i) B. Male bilateral hemispheric variance

An overview observation on bilateral hemispheric variance in males shows a greater tract mean length (mm) observed in MGH_1022 on the right side (152.26), but the left with only 6284.25 tract length mean. Similarly, a least tract mean length (mm) i.e. 85.78 is seen on right side in male subject MGH_1006 and 1873.12 on the left side (Table-19).

**Table 19:**
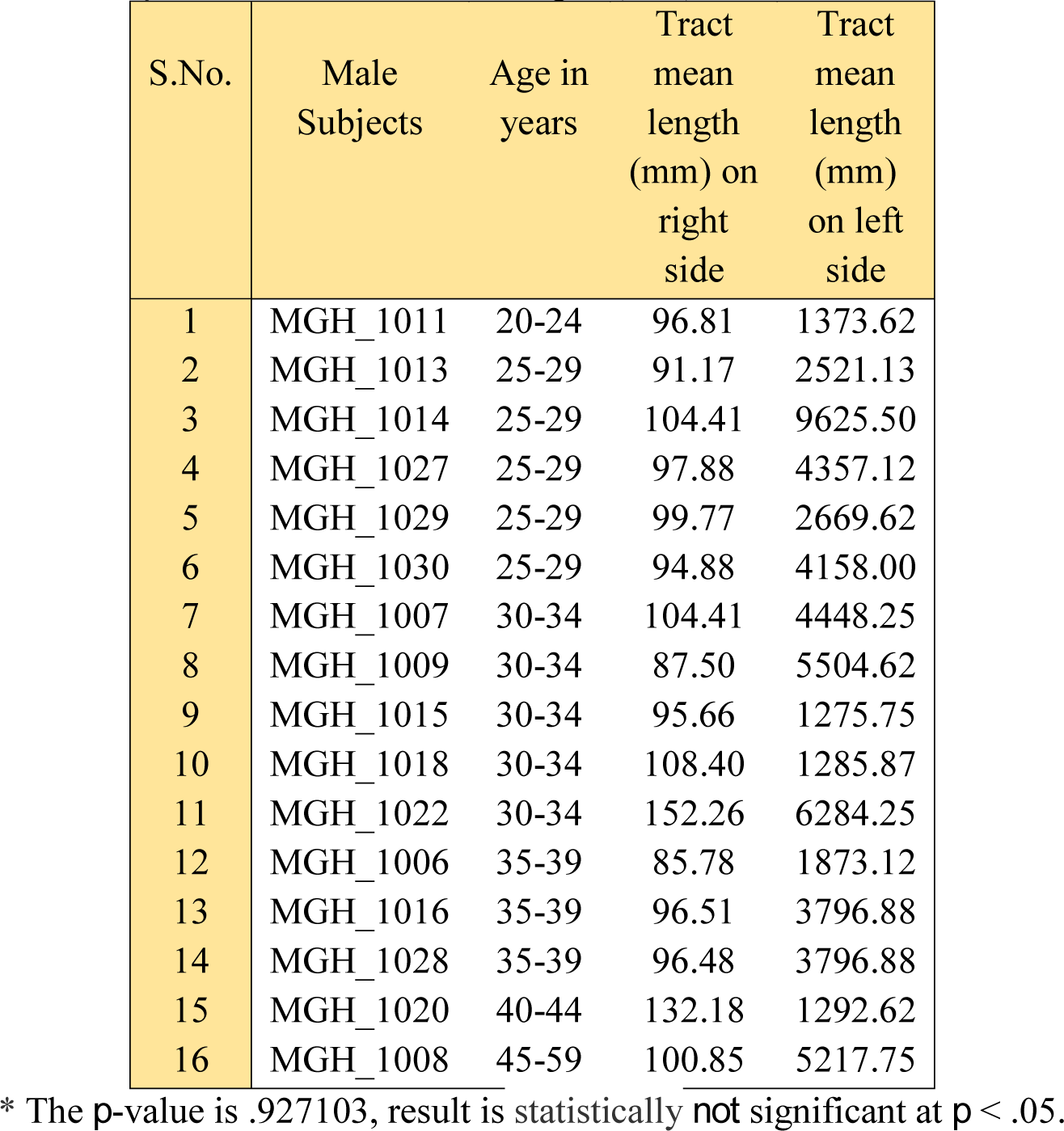
Comparison of Tract mean length (mm) on male bilateral variances *

| S.No. | Male Subjects | Age in years | Tract mean length (mm) on right side | Tract mean length (mm) on left side |
| --- | --- | --- | --- | --- |
| 1 | MGH_1011 | 20-24 | 96.81 | 1373.62 |
| 2 | MGH_1013 | 25-29 | 91.17 | 2521.13 |
| 3 | MGH_1014 | 25-29 | 104.41 | 9625.50 |
| 4 | MGH_1027 | 25-29 | 97.88 | 4357.12 |
| 5 | MGH_1029 | 25-29 | 99.77 | 2669.62 |
| 6 | MGH_1030 | 25-29 | 94.88 | 4158.00 |
| 7 | MGH_1007 | 30-34 | 104.41 | 4448.25 |
| 8 | MGH_1009 | 30-34 | 87.50 | 5504.62 |
| 9 | MGH_1015 | 30-34 | 95.66 | 1275.75 |
| 10 | MGH_1018 | 30-34 | 108.40 | 1285.87 |
| 11 | MGH_1022 | 30-34 | 152.26 | 6284.25 |
| 12 | MGH_1006 | 35-39 | 85.78 | 1873.12 |
| 13 | MGH_1016 | 35-39 | 96.51 | 3796.88 |
| 14 | MGH_1028 | 35-39 | 96.48 | 3796.88 |
| 15 | MGH_1020 | 40-44 | 132.18 | 1292.62 |
| 16 | MGH_1008 | 45-59 | 100.85 | 5217.75 |

\* The $p$ -value is .927103, result is statistically *not* significant at $p < .05$ .

Conclusively, for the set of results observed in male bilateral hemispheric variance, we noted that there is no significant difference in the tract mean length (mm) ending in both hemispheres (Table-19), hence in support of our subsidiary thesis (Graph-9).

**Graph-9:**
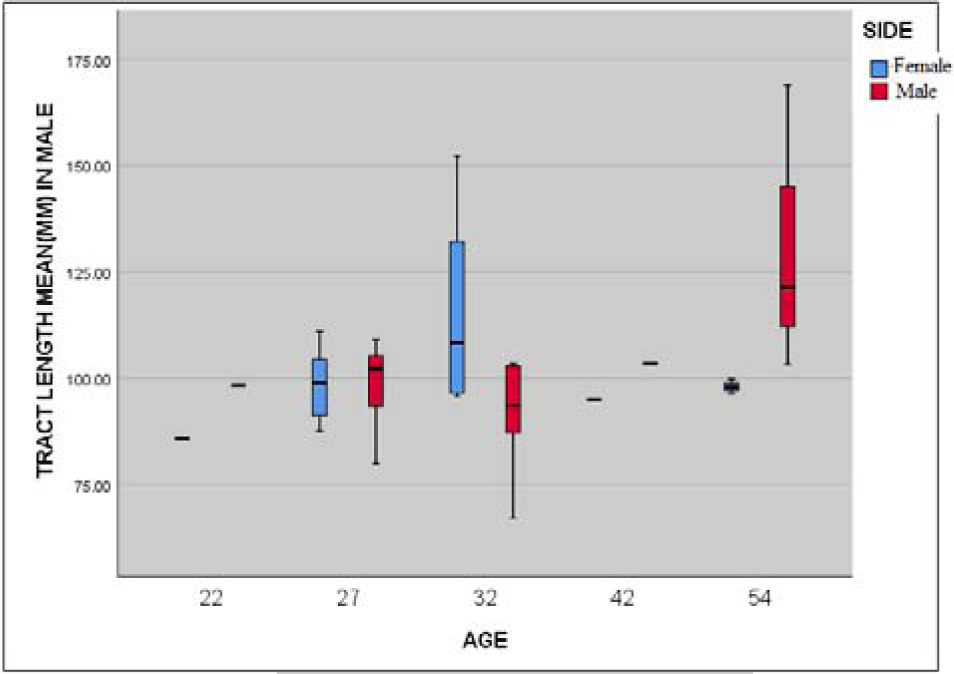
Graphical representation of tract length mean (mm) between left and right sides in male subjects.

#### ii) A. Female unilateral hemispheric variance

Similarly, on right side for the16 female datasets analysed (Table-20), we found that the subject MGH_1021 (Fig-21) with the mean age of 32 years have the highest tract length mean (mm) (150.13), and the dataset MGH_1019 (Fig-22) with the mean age of 37 with least tract length mean (mm) (57.67).

**Fig 21:**
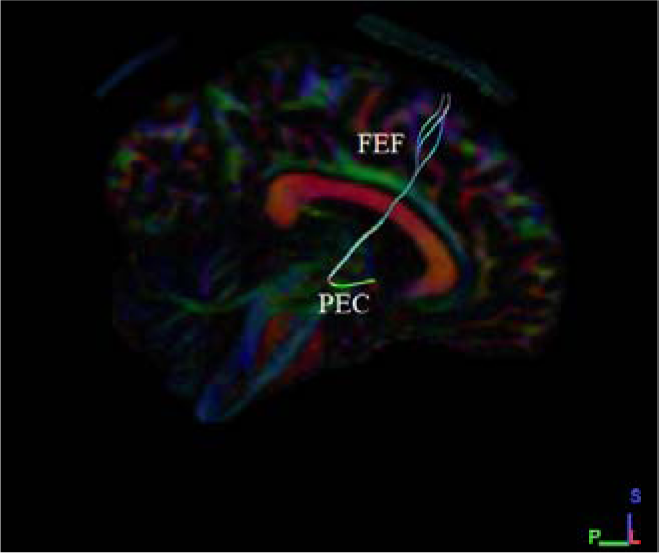
Shows the sagittal section with highest (150.13) tract length mean (mm) in female subject (MGH_1021) on right side.

**Fig 22:**
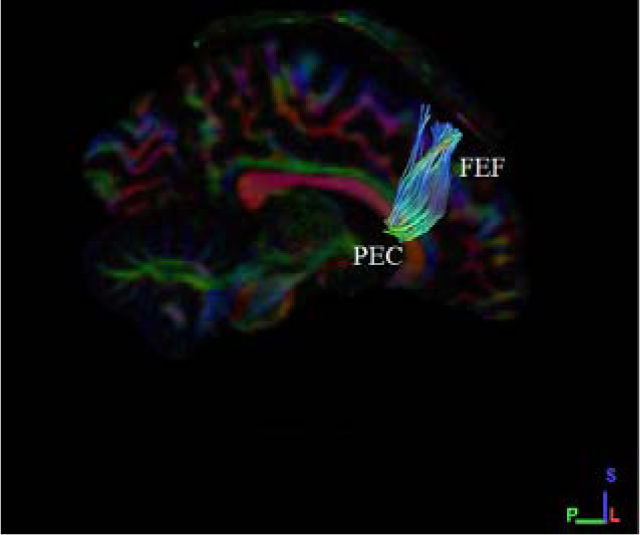
Shows the coronal section with least (56.67) Tract mean length (mm) in female subject (MGH_10019) right on right side.

**Table 20:**
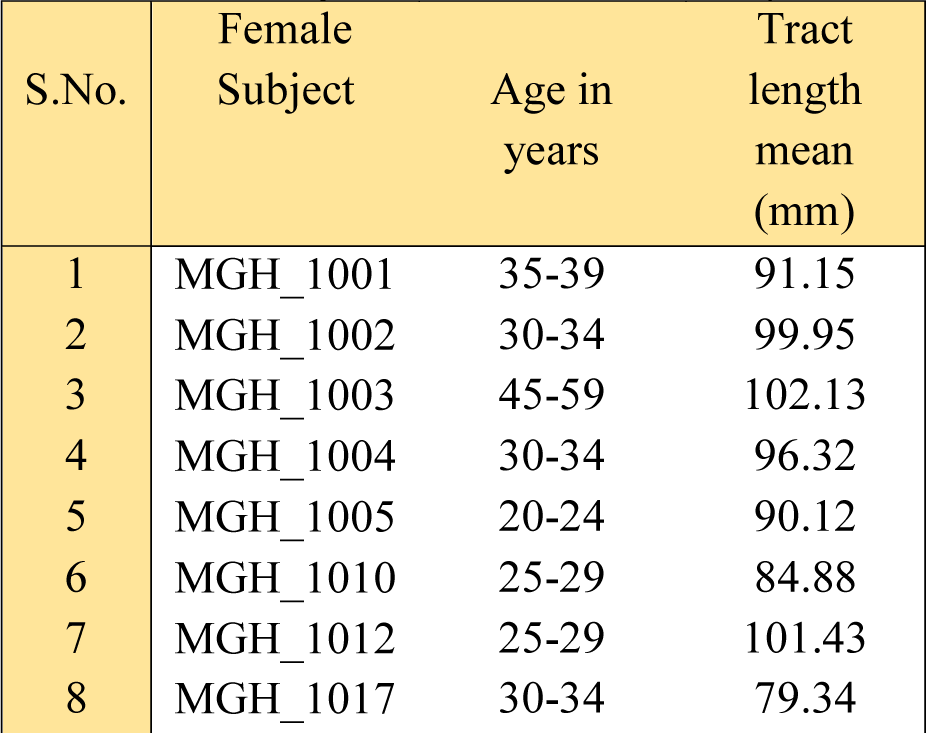
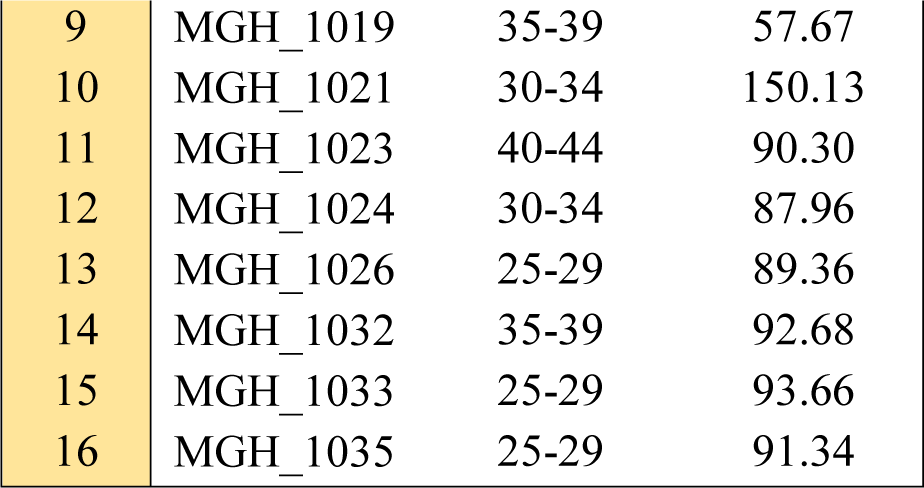
Tract mean length (mm) in female subjects on right side

An observation of 16 female left side datasets (Table-21) showed subject MGH_1021 (Fig-23) with the mean age of 32 years to have highest tract mean length (mm) (198.78), and the dataset MGH_1017 (Fig-24) with the mean age of 32 with least tract mean length (mm) (81.41).

**Fig 23:**
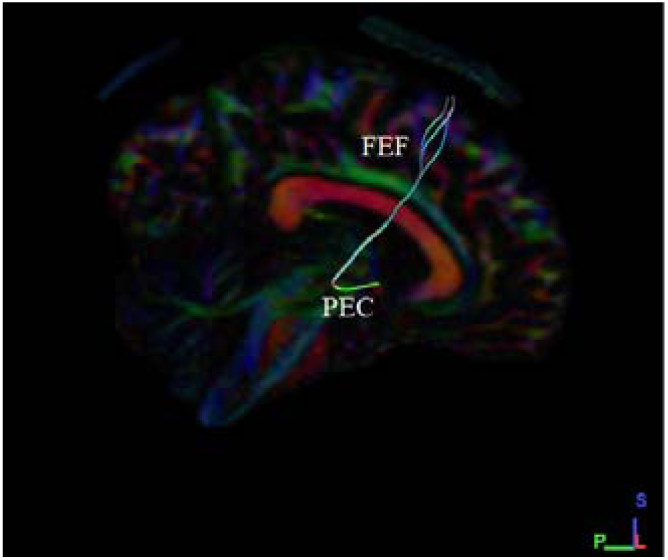
Shows the sagittal section with highest (198.78) tract length mean (mm) in female subject (MGH_1021) on left side.

**Fig 24:**
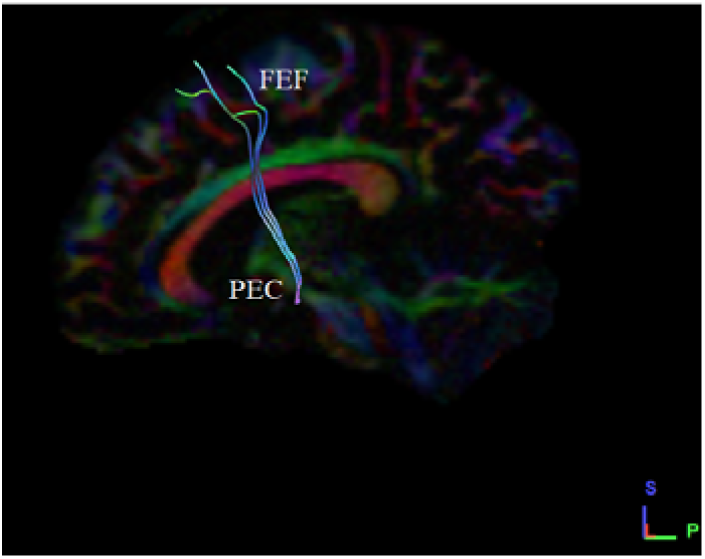
Shows the sagittal section with least (81.41) tract length mean (mm) in female subject (MGH_1017) on left side.

**Table 21:** Tract mean length (mm) in female subjects on left side

| S.No. | Female Subject | Age in years | Tract length mean (mm) |
| --- | --- | --- | --- |
| 1 | MGH_1001 | 35-39 | 93.08 |
| 2 | MGH_1002 | 30-34 | 91.53 |
| 3 | MGH_1003 | 45-59 | 102.11 |
| 4 | MGH_1004 | 30-34 | 97.40 |
| 5 | MGH_1005 | 20-24 | 111.07 |
| 6 | MGH_1010 | 25-29 | 98.12 |
| 7 | MGH_1012 | 25-29 | 106.31 |
| 8 | MGH_1017 | 30-34 | 81.41 |
| 9 | MGH_1019 | 35-39 | 84.20 |
| 10 | MGH_1021 | 30-34 | 198.78 |
| 11 | MGH_1023 | 40-44 | 102.55 |
| 12 | MGH_1024 | 30-34 | 93.26 |
| 13 | MGH_1026 | 25-29 | 94.61 |
| 14 | MGH_1032 | 35-39 | 99.31 |
| 15 | MGH_1033 | 25-29 | 97.38 |
| 16 | MGH_1035 | 25-29 | 94.11 |

#### ii) B. Female bilateral hemispheric variance

An overview observation on bilateral hemispheric variances in female shows a greater tract length mean (mm) observed in MGH_1021 on the right side (150.13) with the mean age of 32 years, but the left with only 198.78 tract length mean (mm). Similarly, a least tract length mean (mm) i.e. 57.67 is seen on right side in subject MGH_1019 and 84.20 on the left side (Table-22).

**Table 22:**
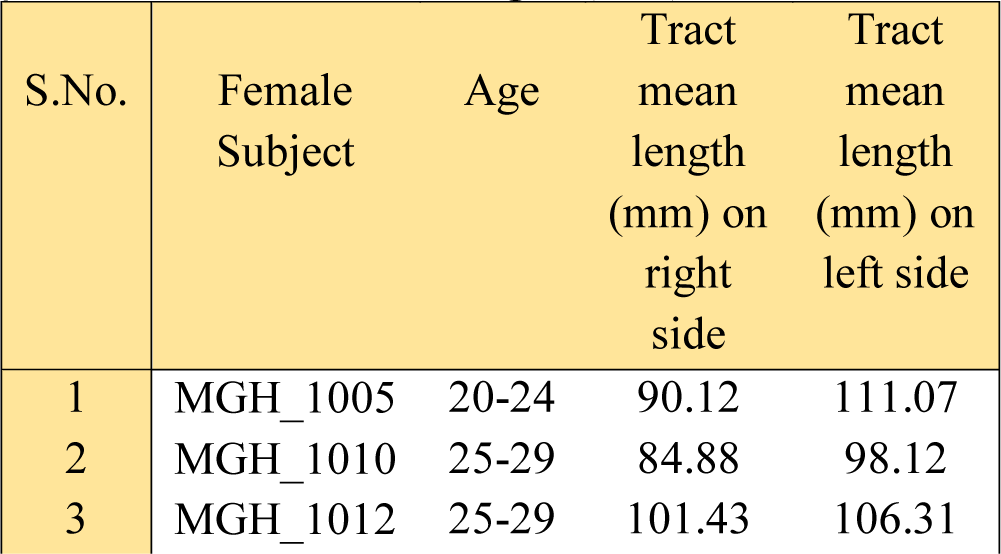
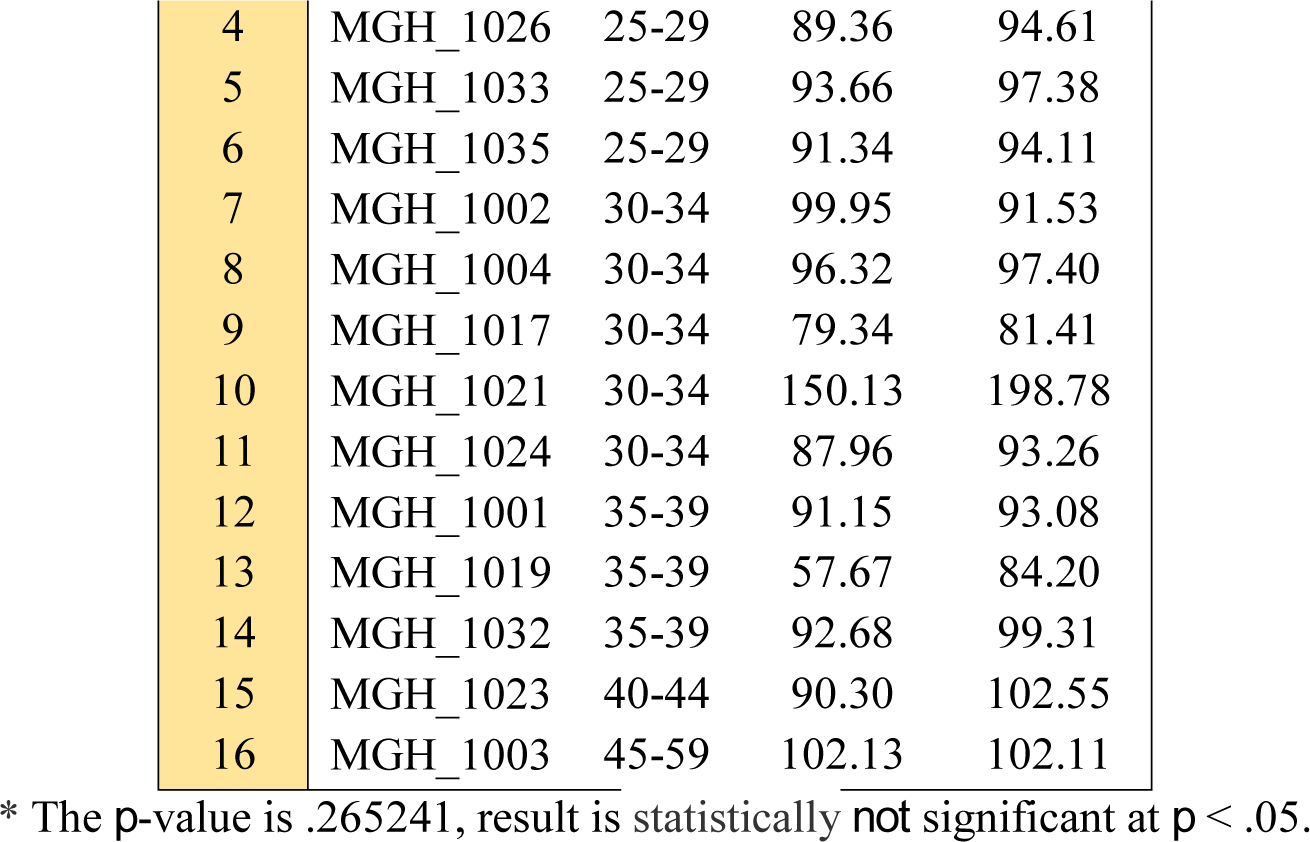
Comparison of Tract mean length (mm) on female bilateral variances *

Conclusively, for the set of results observed in female bilateral hemispheric variance, we noted no difference exists in the tract mean length (mm) ending in the two hemispheres (Table-22) upholding our subsidiary hypothesis (Graph-10).

**Graph-10:**
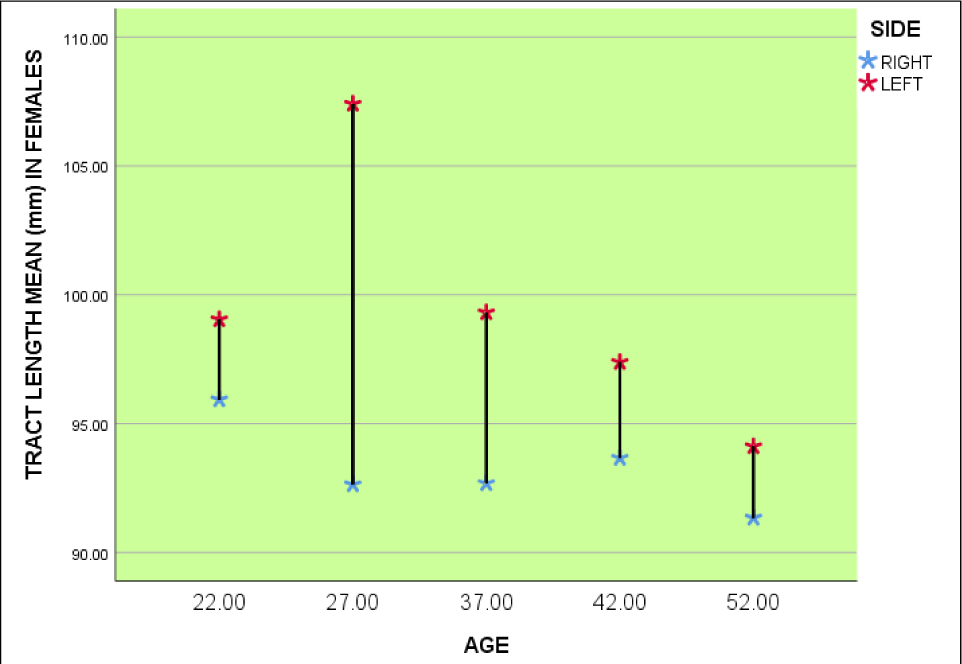
Graphical representation of tract length mean (mm) between left and right sides in female subjects.

#### iii) Male and female unilateral hemispheric variances (right side)

On right side for the 16 male datasets analysed (Table-17), we found that the subject MGH_1022 (Fig-17) with the mean age of 42 years have the highest tract length mean (mm) (152.26), and the datasets MGH_1006 (Fig-18) with the mean age of 37 with least tract length mean (mm) (85.78).

Similarly, on right side for the 16 female datasets analysed (Table-20), we found that the subject MGH_1021 (Fig-21) with the mean age of 32 years have the highest tract length mean (mm) (150.13), and the dataset MGH_1019 (Fig-22) with the mean age of 37 with least tract length mean (mm) (57.67).

Conclusively, for the set of results observed in male and female unilateral hemispheric variances on right side, we noted no difference exists in the tract length mean (mm) in the hemispheres (Table-23), results observed were similar in both male and female unilateral hemispheres (Graph-11), upholding our secondary conjecture.

**Table 23:** Comparison between Tract mean length (mm) of both male and female unilateral hemispheric variances on right side *

| S.No. | Tract length mean (mm) on right side in male subjects | Tract length mean (mm) on right side in female subjects |
| --- | --- | --- |
| 1 | 96.81 | 90.12 |
| 2 | 91.17 | 84.88 |
| 3 | 104.41 | 101.43 |
| 4 | 97.88 | 89.36 |
| 5 | 99.77 | 93.66 |
| 6 | 94.88 | 91.34 |
| 7 | 104.41 | 99.95 |
| 8 | 87.50 | 96.32 |
| 9 | 95.66 | 79.34 |
| 10 | 108.40 | 150.13 |
| 11 | 152.26 | 87.96 |
| 12 | 85.78 | 91.15 |
| 13 | 96.51 | 57.67 |
| 14 | 96.48 | 92.68 |
| 15 | 132.18 | 90.30 |
| 16 | 100.85 | 102.13 |

\* The $p$ -value is .135945, result is statistically not significant at $p < .05$ .

Correlative observation on unilateral hemispheric variances in male and female on right side is shown in Table-23.

#### iv) Male and female unilateral hemispheric variances (left side)

An observation of left side of the 16 male datasets (Table-18) showed subject MGH_1014 (Fig-19) with the mean age of 27 years have the highest tract mean length (mm) (9625.50), and the dataset MGH_1015 (Fig-20) with the mean age of 32 with least tract mean length (mm) (1275.75).

An observation of left side of the 16 female datasets (Table-21) showed subject MGH_1021 (Fig-23) with the mean age of 32 years having the highest tract mean length (mm) (198.78), and the dataset MGH_1017 (Fig-24) with the mean age of 32 with least tract mean length (mm) (81.41).

Conclusively, for our set of results of male and female unilateral hemispheric variances on left side, we noted no difference exists in the tract length mean (mm) in the hemispheres (Table-22), results observed were similar in both male and female unilateral hemispheres (Graph-12), hence in support of our secondary hypothesis.

Correlative observation on unilateral hemispheric variances in male and female subjects on left side is shown in Table-24.

**Table 24:** Comparison between Tract mean length (mm) of both male and female unilateral hemispheric variances on right side *

| S.No. | Tract length mean (mm) on left side in male subjects | Tract length mean (mm) on left side in female subjects |
| --- | --- | --- |
| 1 | 1373.62 | 111.07 |
| 2 | 2521.13 | 98.12 |
| 3 | 9625.50 | 106.31 |
| 4 | 4357.12 | 94.61 |
| 5 | 2669.62 | 97.38 |
| 6 | 4158.00 | 94.11 |
| 7 | 4448.25 | 91.53 |
| 8 | 5504.62 | 97.40 |
| 9 | 1275.75 | 81.41 |
| 10 | 1285.87 | 198.78 |
| 11 | 6284.25 | 93.26 |
| 12 | 1873.12 | 93.08 |
| 13 | 3796.88 | 84.20 |
| 14 | 3796.88 | 99.31 |
| 15 | 1292.62 | 102.55 |
| 16 | 5217.75 | 102.11 |

\* The $p$ -value is .978036, result is statistically not significant at $p < .05$ .

**Graph-11:**
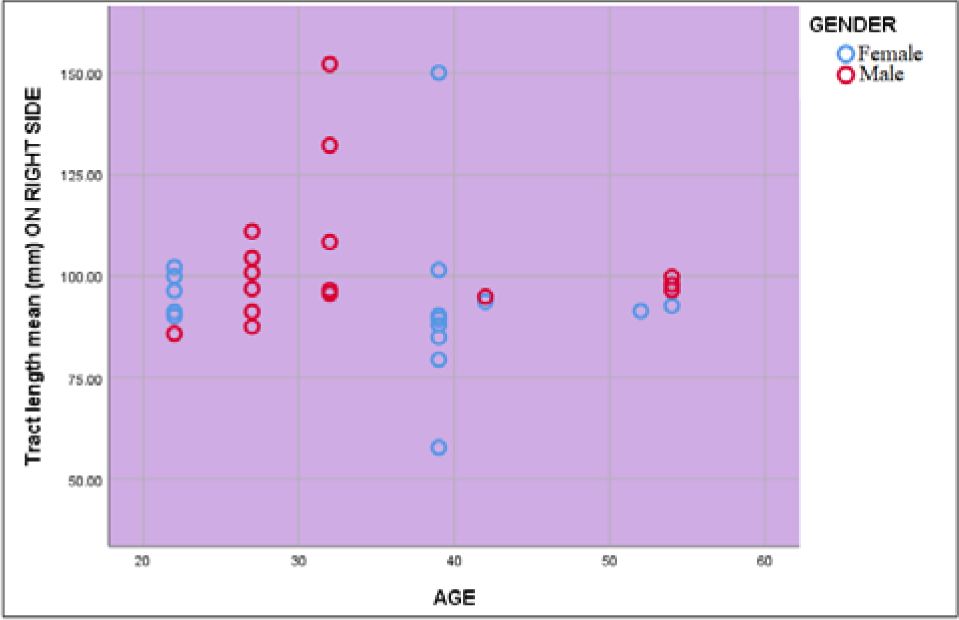
Graphical representation of tract length mean (mm) on male and female subjects on right side.

**Graph-12:**
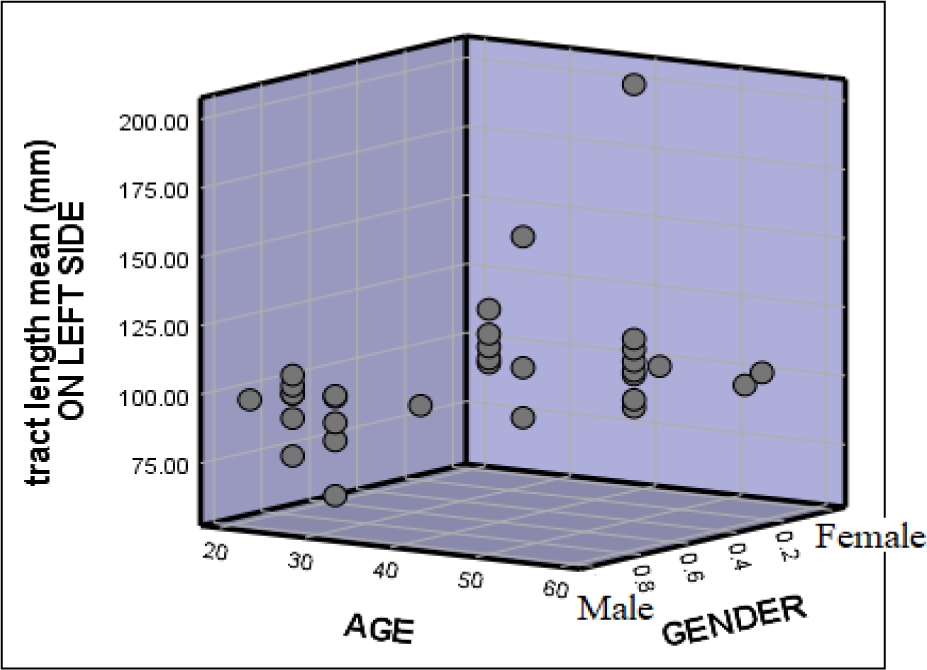
Graphical representation of tract length mean (mm) on male and female subjects on left side.

### D) Tract length standard deviation (mm)

Tract length standard deviation is a measure that is used to quantify the amount of variation or dispersion a set of data values exhibit. We analyzed tract mean length (mm) present in both the sexes i.e. i) male subjects and ii) female subjects.

#### i) A. Male unilateral hemispheric variance

On right side for the 16 male datasets analysed (Table-25), we found that the subject MGH_1022 (Fig-25) with the mean age of 32 years have the highest tract length standard deviation (mm) (152.26), and the dataset MGH_1028 (Fig-26) with the mean age of 37 with least tract length mean (mm) (85.78).

**Fig 25:**
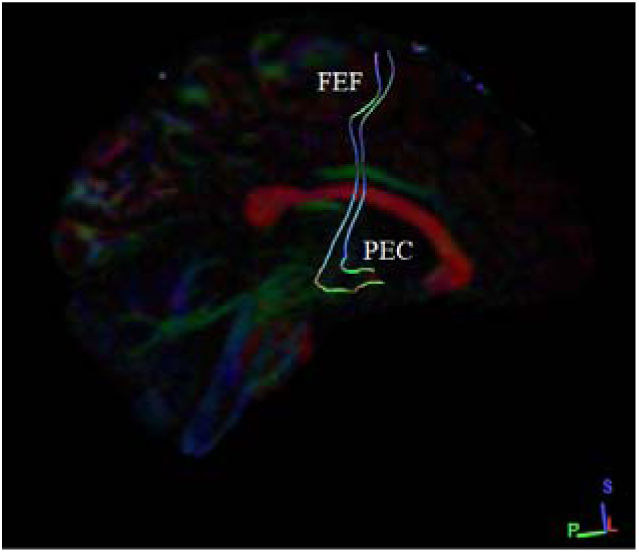
Shows the sagittal section with highest (61.50) Tract length standard deviation (mm) in male subject (MGH_1022) on right side.

**Fig 26:**
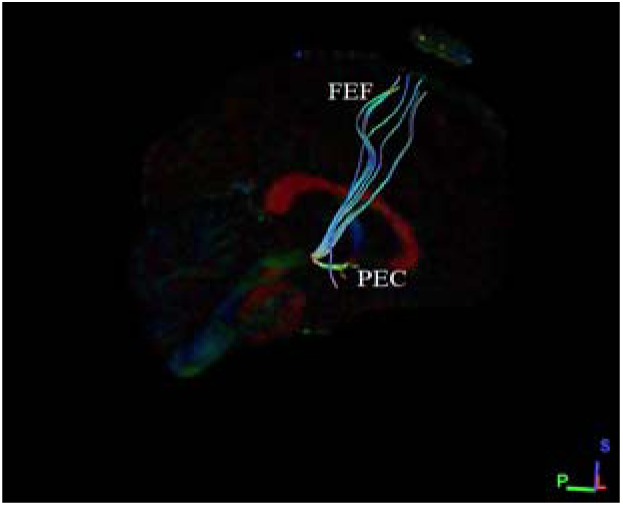
Shows the sagittal section with least (2.41) Tract length standard deviation (mm) in male subject (MGH_1028) on right side.

**Table 25:** Tract length standard deviation (mm) in male subjects on right side

| S.No. | Male Subjects | Age in years | Tract length standard deviation (mm) |
| --- | --- | --- | --- |
| 1 | MGH_1006 | 35-39 | 13.84 |
| 2 | MGH_1007 | 30-34 | 16.84 |
| 3 | MGH_1008 | 45-59 | 3.98 |
| 4 | MGH_1009 | 30-34 | 7.63 |
| 5 | MGH_1011 | 20-24 | 4.06 |
| 6 | MGH_1013 | 25-29 | 9.36 |
| 7 | MGH_1014 | 25-29 | 12.85 |
| 8 | MGH_1015 | 30-34 | 19.64 |
| 9 | MGH_1016 | 35-39 | 7.36 |
| 10 | MGH_1018 | 30-34 | 27.02 |
| 11 | MGH_1020 | 40-44 | 5.16 |
| 12 | MGH_1022 | 30-34 | 61.50 |
| 13 | MGH_1027 | 25-29 | 11.10 |
| 14 | MGH_1028 | 35-39 | 2.41 |
| 15 | MGH_1029 | 25-29 | 4.75 |
| 16 | MGH_1030 | 25-29 | 5.39 |

On left side for the 16 male datasets analysed (Table-26), we found that the subject MGH_1028 (Fig-27) with the mean age of 37 years have highest tract length standard deviation (mm) (79.59), and the dataset MGH_1018 (Fig-28) with the mean age of 32 with least tract length mean (mm) (2.95).

**Fig 27:**
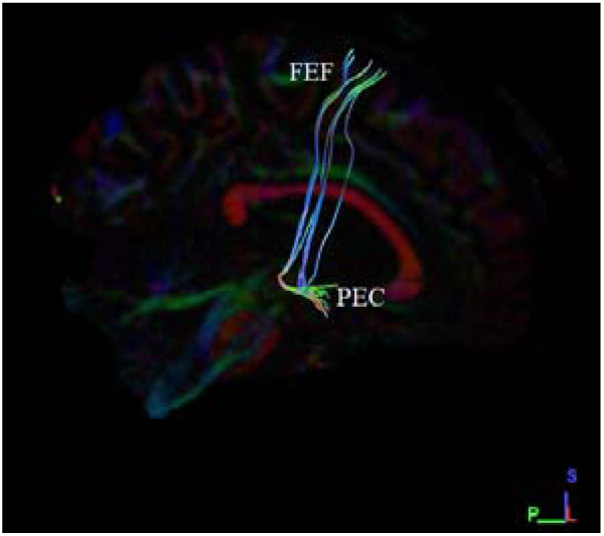
Shows the sagittal section with highest (79.59) tract length standard deviation (mm) in male subject (MGH_1028) on left side.

**Fig 28:**
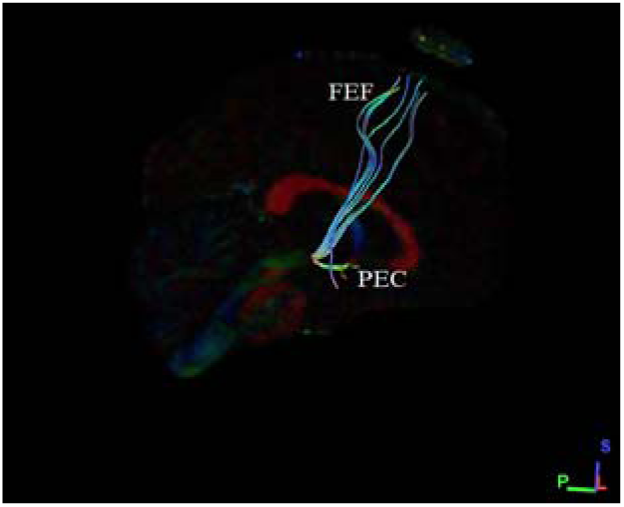
Shows the sagittal section with least (2.95) tract length standard deviation (mm) in male subject (MGH_1018) on left side.

**Table 26:**
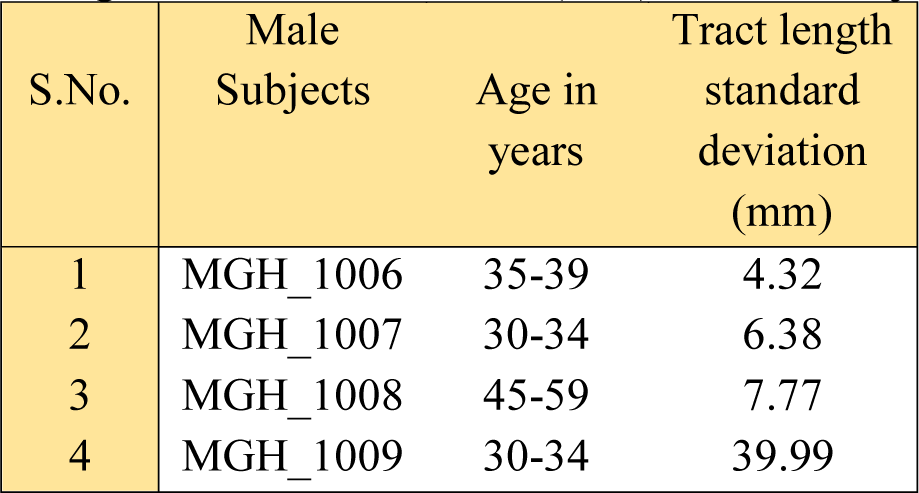
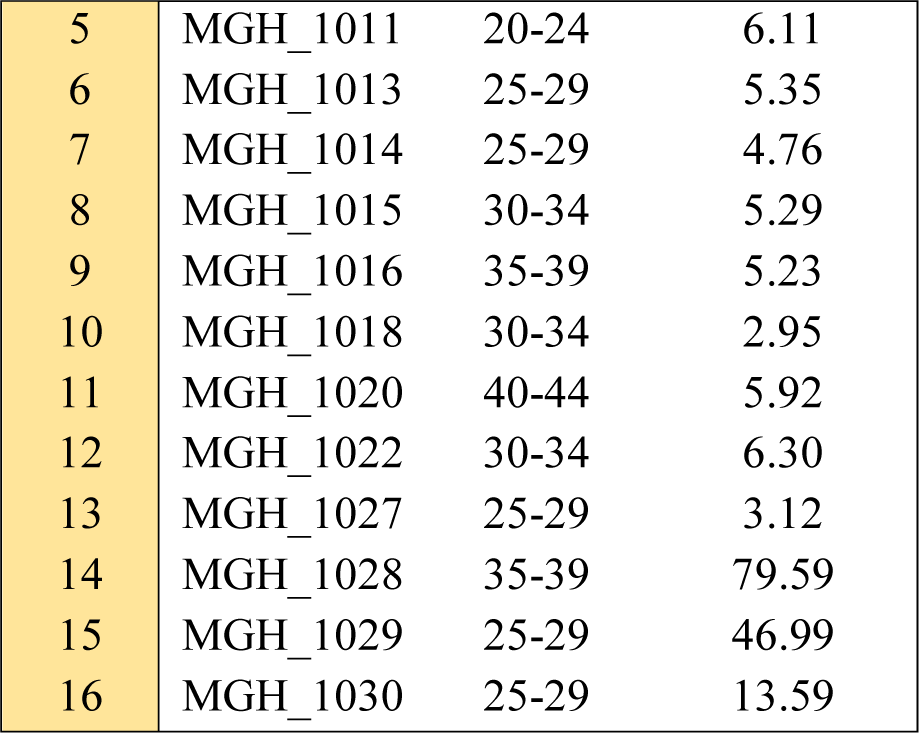
Tract length standard deviation (mm) in male subjects on left side

| S.No. | Male Subjects | Age in years | Tract length standard deviation (mm) |
| --- | --- | --- | --- |
| 1 | MGH_1006 | 35-39 | 4.32 |
| 2 | MGH_1007 | 30-34 | 6.38 |
| 3 | MGH_1008 | 45-59 | 7.77 |
| 4 | MGH_1009 | 30-34 | 39.99 |
| 5 | MGH_1011 | 20-24 | 6.11 |
| 6 | MGH_1013 | 25-29 | 5.35 |
| 7 | MGH_1014 | 25-29 | 4.76 |
| 8 | MGH_1015 | 30-34 | 5.29 |
| 9 | MGH_1016 | 35-39 | 5.23 |
| 10 | MGH_1018 | 30-34 | 2.95 |
| 11 | MGH_1020 | 40-44 | 5.92 |
| 12 | MGH_1022 | 30-34 | 6.30 |
| 13 | MGH_1027 | 25-29 | 3.12 |
| 14 | MGH_1028 | 35-39 | 79.59 |
| 15 | MGH_1029 | 25-29 | 46.99 |
| 16 | MGH_1030 | 25-29 | 13.59 |

#### i) B. Male bilateral hemispheric variance

An overview observation on bilateral hemispheric variance in males shows a greater tract length standard deviation (mm) observed in MGH_1022 on the right side (61.50), but the left with only 6.30 tract length standard deviation (mm). Similarly, a least tract length standard deviation (mm) i.e. 2.41 is seen on right side in male subject MGH_1028 and 79.59 on the left side (Table-27).

**Table 27:** Comparison of Tract length standard deviation (mm) on male bilateral variances *

| S.No. | Male Subjects | Age in years | Tract length standard deviation (mm) on right side | Tract length standard deviation (mm) on left side |
| --- | --- | --- | --- | --- |
| 1 | MGH_1011 | 20-24 | 4.06 | 6.11 |
| 2 | MGH_1013 | 25-29 | 9.36 | 5.35 |
| 3 | MGH_1014 | 25-29 | 12.85 | 4.76 |
| 4 | MGH_1027 | 25-29 | 11.10 | 3.12 |
| 5 | MGH_1029 | 25-29 | 4.75 | 46.99 |
| 6 | MGH_1030 | 25-29 | 5.39 | 13.59 |
| 7 | MGH_1007 | 30-34 | 16.84 | 6.38 |
| 8 | MGH_1009 | 30-34 | 7.63 | 39.99 |
| 9 | MGH_1015 | 30-34 | 19.64 | 5.29 |
| 10 | MGH_1018 | 30-34 | 27.02 | 2.95 |
| 11 | MGH_1022 | 30-34 | 61.50 | 6.30 |
| 12 | MGH_1006 | 35-39 | 7.36 | 4.32 |
| 13 | MGH_1016 | 35-39 | 7.36 | 5.23 |
| 14 | MGH_1028 | 35-39 | 2.41 | 79.59 |
| 15 | MGH_1020 | 40-44 | 5.16 | 5.92 |
| 16 | MGH_1008 | 45-59 | 3.98 | 7.77 |

\* The p-value is .769128, result is statistically *not* significant at $p < .05$ .

Conclusively, for our set of results of male bilateral hemispheric variance, we noted that no difference exists in the tract length standard deviation (mm) ending in the two hemispheres (Table-27), hence upholding our secondary hypothesis (Graph-13).

**Graph-13:**
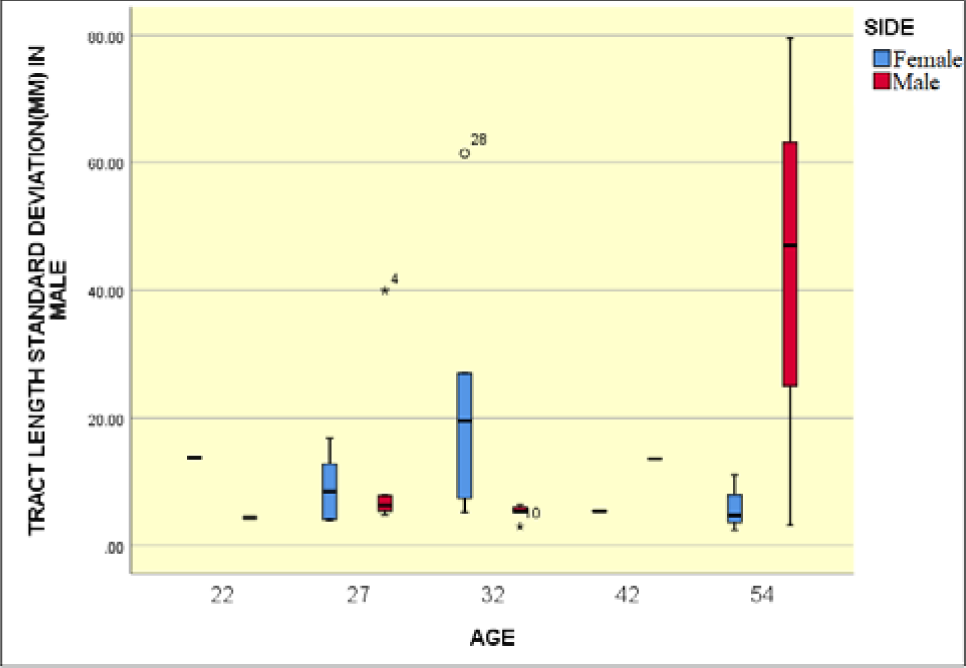
Graphical representation of tract length standard deviation (mm) between left and right sides in male subjects.

#### ii) A. Female unilateral hemispheric variance

Similarly, on right side for the16 female datasets analysed (Table-28), we found that the subject MGH_1021 (Fig-29) with the mean age of 32 years have the highest tract length standard deviation (mm) (64.38), and the dataset MGH_1019 (Fig-30) with the mean age of 37 have the least tract length standard deviation (mm) (1.21).

**Fig 29:**
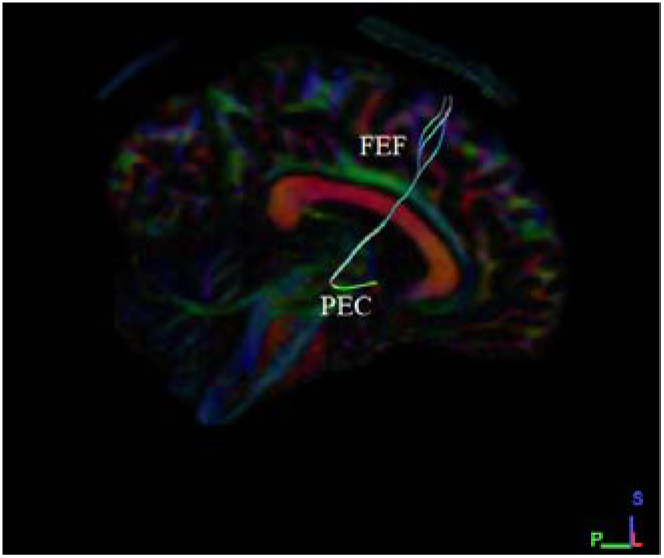
Shows the sagittal section with highest (64.38) tract length standard deviation (mm) in female subject (MGH_1021) on right side.

**Fig 30:**
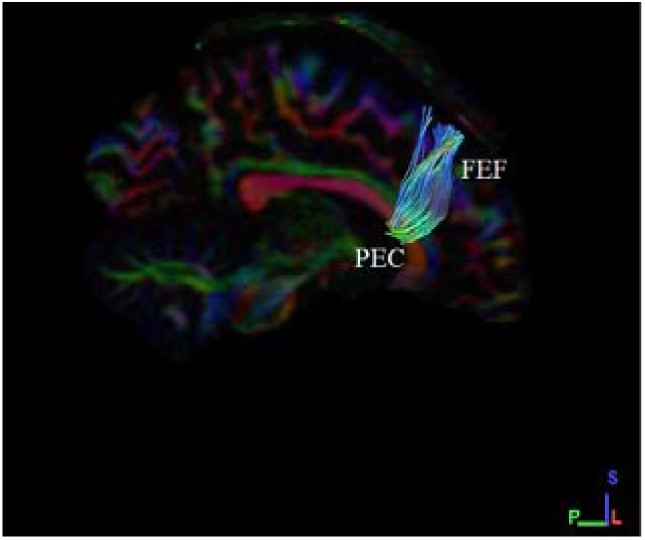
Shows the sagittal section with least (1.21) tract length standard deviation (mm) in female subject (MGH_1019) on right side.

**Table 28:** Tract length standard deviation (mm) in female subjects on right side

| S.No. | Female Subject | Age in years | Tract length standard deviation (mm) |
| --- | --- | --- | --- |
| 1 | MGH_1001 | 35-39 | 17.64 |
| 2 | MGH_1002 | 30-34 | 22.64 |
| 3 | MGH_1003 | 45-59 | 8.07 |
| 4 | MGH_1004 | 30-34 | 5.87 |
| 5 | MGH_1005 | 20-24 | 10.50 |
| 6 | MGH_1010 | 25-29 | 11.38 |
| 7 | MGH_1012 | 25-29 | 7.69 |
| 8 | MGH_1017 | 30-34 | 8.13 |
| 9 | MGH_1019 | 35-39 | 1.21 |
| 10 | MGH_1021 | 30-34 | 64.38 |
| 11 | MGH_1023 | 40-44 | 15.12 |
| 12 | MGH_1024 | 30-34 | 6.75 |
| 13 | MGH_1026 | 25-29 | 8.52 |
| 14 | MGH_1032 | 35-39 | 1.51 |
| 15 | MGH_1033 | 25-29 | 9.35 |
| 16 | MGH_1035 | 25-29 | 8.87 |

An observation of the 16 female left side datasets (Table-29) showed the subject MGH_1019 (Fig-31) with the mean age of 37 years having the highest length standard deviation (mm) (77.31), and the dataset MGH_1012 (Fig-32) with the mean age of 27 with least length standard deviation (mm) (2.49).

**Fig 31:**
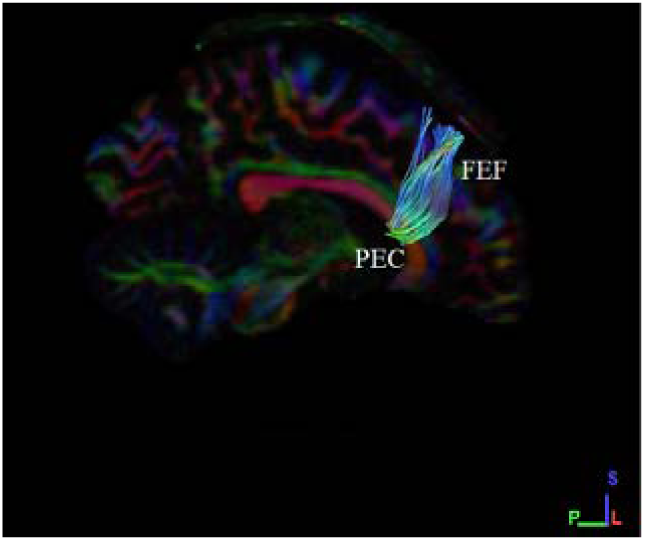
Shows the sagittal section with highest tract length standard deviation (mm) in female subject (MGH_1019) on left side.

**Fig 32:**
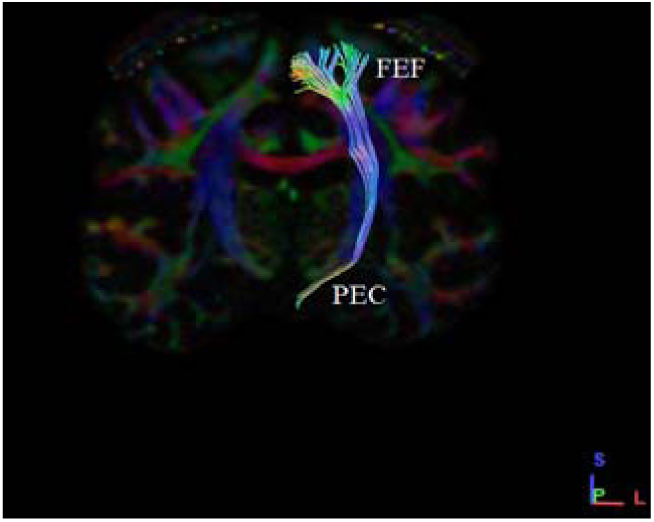
Shows the coronal section with least tract length standard deviation (mm) in female subject (MGH_1012) on left side.

**Table 29:**
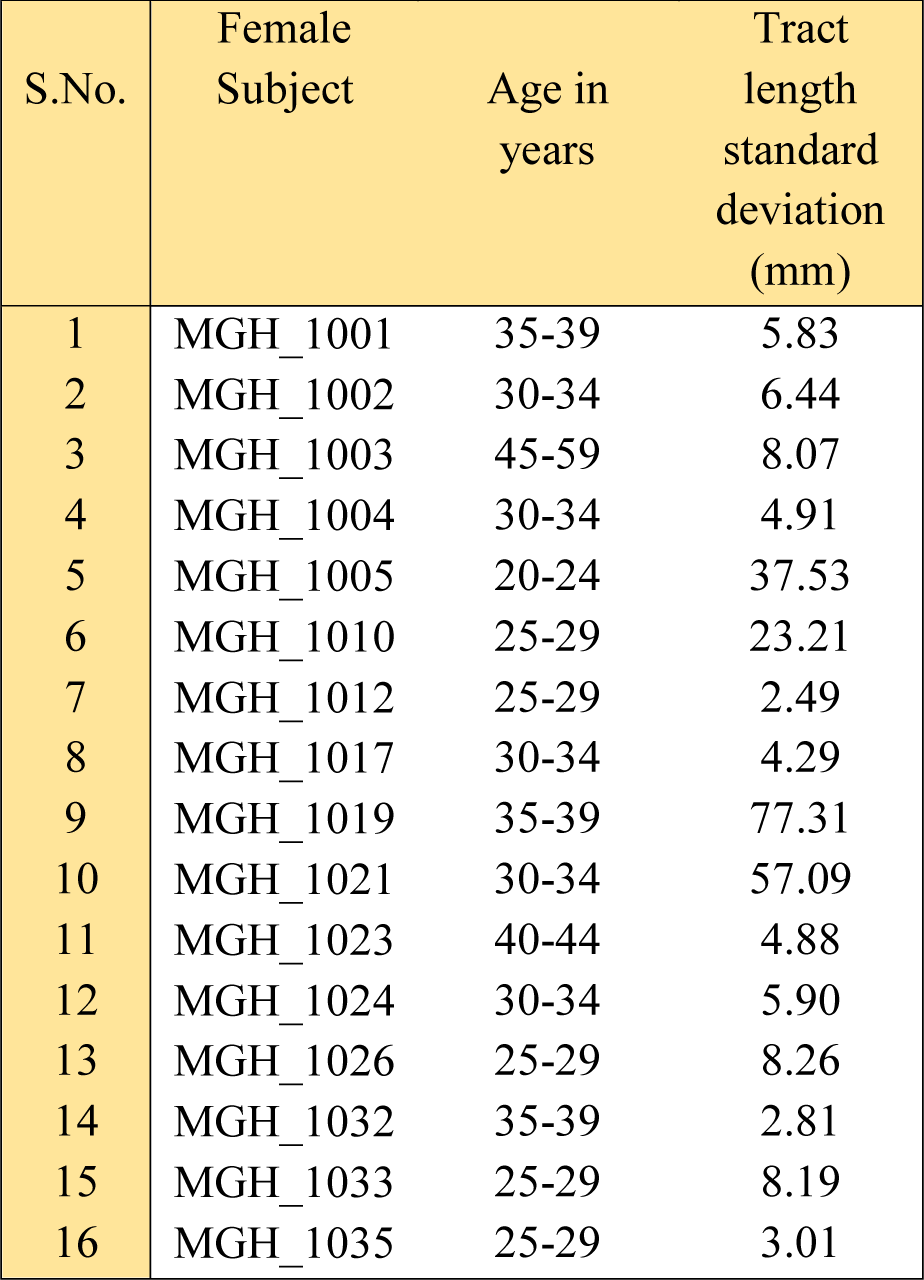
Tract length standard deviation (mm) in female subjects on left side

| S.No. | Female Subject | Age in years | Tract length standard deviation (mm) |
| --- | --- | --- | --- |
| 1 | MGH_1001 | 35-39 | 5.83 |
| 2 | MGH_1002 | 30-34 | 6.44 |
| 3 | MGH_1003 | 45-59 | 8.07 |
| 4 | MGH_1004 | 30-34 | 4.91 |
| 5 | MGH_1005 | 20-24 | 37.53 |
| 6 | MGH_1010 | 25-29 | 23.21 |
| 7 | MGH_1012 | 25-29 | 2.49 |
| 8 | MGH_1017 | 30-34 | 4.29 |
| 9 | MGH_1019 | 35-39 | 77.31 |
| 10 | MGH_1021 | 30-34 | 57.09 |
| 11 | MGH_1023 | 40-44 | 4.88 |
| 12 | MGH_1024 | 30-34 | 5.90 |
| 13 | MGH_1026 | 25-29 | 8.26 |
| 14 | MGH_1032 | 35-39 | 2.81 |
| 15 | MGH_1033 | 25-29 | 8.19 |
| 16 | MGH_1035 | 25-29 | 3.01 |

#### ii) B. Female bilateral hemispheric variance

An overview observation on bilateral hemispheric variance in female shows a greater tract length standard deviation (mm) observed in MGH_1021 on the right side (150.13) with the mean age of 32 years, but the left with only 198. Similarly, a least tract length standard deviation (mm) i.e. 57.67 is seen on right side in subject MGH_1019 and 84.20 on the left side (Table-30).

**Table 30:** Comparison of Tract length standard deviation (mm) on female bilateral variances *

| S.No. | Female Subject | Age | Tract length standard deviation (mm) on right side | Tract length standard deviation (mm) on left side |
| --- | --- | --- | --- | --- |
| 1 | MGH_1005 | 20-24 | 10.50 | 37.53 |
| 2 | MGH_1010 | 25-29 | 11.38 | 23.21 |
| 3 | MGH_1012 | 25-29 | 7.69 | 2.49 |
| 4 | MGH_1026 | 25-29 | 8.52 | 8.26 |
| 5 | MGH_1033 | 25-29 | 9.35 | 8.19 |
| 6 | MGH_1035 | 25-29 | 8.87 | 3.01 |
| 7 | MGH_1002 | 30-34 | 22.64 | 6.44 |
| 8 | MGH_1004 | 30-34 | 5.87 | 4.91 |
| 9 | MGH_1017 | 30-34 | 8.13 | 4.29 |
| 10 | MGH_1021 | 30-34 | 64.38 | 57.09 |
| 11 | MGH_1024 | 30-34 | 6.75 | 5.90 |
| 12 | MGH_1001 | 35-39 | 17.64 | 5.83 |
| 13 | MGH_1019 | 35-39 | 1.21 | 77.31 |
| 14 | MGH_1032 | 35-39 | 1.51 | 2.81 |
| 15 | MGH_1023 | 40-44 | 15.12 | 4.88 |
| 16 | MGH_1003 | 45-59 | 22.64 | 3.01 |

\* The $p$ -value is .624608, result is statistically *not* significant at $p < .05$ .

Conclusively, for our set of results of female bilateral hemispheric variance, we noted no difference exists in the tract length standard deviation (mm) ending in the two hemispheres (Table-30), hence upholding our secondary conjecture.

**Graph-14:**
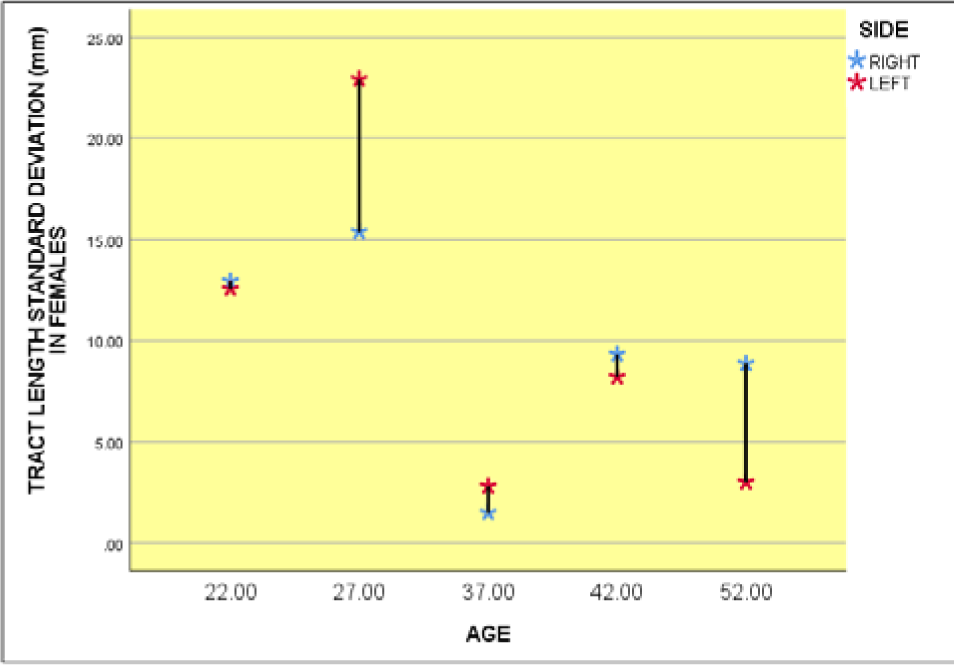
Graphical representation of tract length standard deviation (mm) between left and right sides in female subjects.

#### iii) Male and female unilateral hemispheric variances (right side)

On right side for the 16 male datasets analysed (Table-25), we found that the subject MGH_1022 (Fig-25) with the mean age of 32 years have the highest tract length standard deviation (mm) (152.26), and the dataset MGH_1028 (Fig-26) with the mean age of 37 with least tract length mean (mm) (85.78).

Similarly, on right side for the 16 female datasets analysed (Table-28), we found that the subject MGH_1021 (Fig-29) with the mean age of 32 years have the highest tract length standard deviation (mm) (64.38), and the dataset MGH_1019 (Fig-30) with the mean age of 37 with least tract length standard deviation (mm) (1.21).

Conclusively, for our set of results of male and female unilateral hemispheric variances on right side, we noted no significant difference exists in the tract length standard deviation (mm) between the hemispheres (Table-31), almost similar results were observed in both male and female unilateral hemispheres (Graph-15), hence this upholds our subsidiary conjecture.

**Table-31:** Comparison between Tract length standard deviation (mm) of both male and female unilateral hemispheric variances on right side *

| S.No. | Tract length standard deviation (mm) on right side in male subjects | Tract length standard deviation (mm) on right side in female subjects |
| --- | --- | --- |
| 1 | 4.06 | 10.50 |
| 2 | 9.36 | 11.38 |
| 3 | 12.85 | 7.69 |
| 4 | 11.10 | 8.52 |
| 5 | 4.75 | 9.35 |
| 6 | 5.39 | 8.87 |
| 7 | 16.84 | 22.64 |
| 8 | 7.63 | 5.87 |
| 9 | 19.64 | 8.13 |
| 10 | 27.02 | 64.38 |
| 11 | 61.50 | 6.75 |
| 12 | 7.36 | 17.64 |
| 13 | 7.36 | 1.21 |
| 14 | 2.41 | 1.51 |
| 15 | 5.16 | 15.12 |
| 16 | 3.98 | 22.64 |

\* The p-value is .949664, result is statistically not significant at $p < .05$ .

Correlative observation on unilateral hemispheric variances in male and female subjects on right side is shown in Table-31.

#### iv) Male and female unilateral hemispheric variances (left side)

On left side for the 16 male datasets analysed (Table-26), we found that the subject MGH_1028 (Fig-27) with the mean age of 37 years have the highest tract length standard deviation (mm) (79.59), and the dataset MGH_1018 (Fig-28) with the mean age of 32 with least tract length mean (mm) (2.95).

Observation of left side of the 16 female datasets (Table-29) showed that the subject MGH_1019 (Fig-31) with the mean age of 37 years have the highest length standard deviation (mm) (77.31), and the dataset MGH_1012 (Fig-32) with the mean age of 27 with least length standard deviation (mm) (2.49).

Conclusively, for our set of results of male and female unilateral hemispheric variances on left side, we noted no drastic difference exists in the tract length standard deviation (mm) between the hemispheres (Table-32), similar results were observed in both male and female unilateral hemispheres (Graph-16) hence upholding our secondary hypothesis.

Correlative observation on unilateral hemispheric variances in male and female subjects on left side is shown in Table-32.

**Graph-15:**
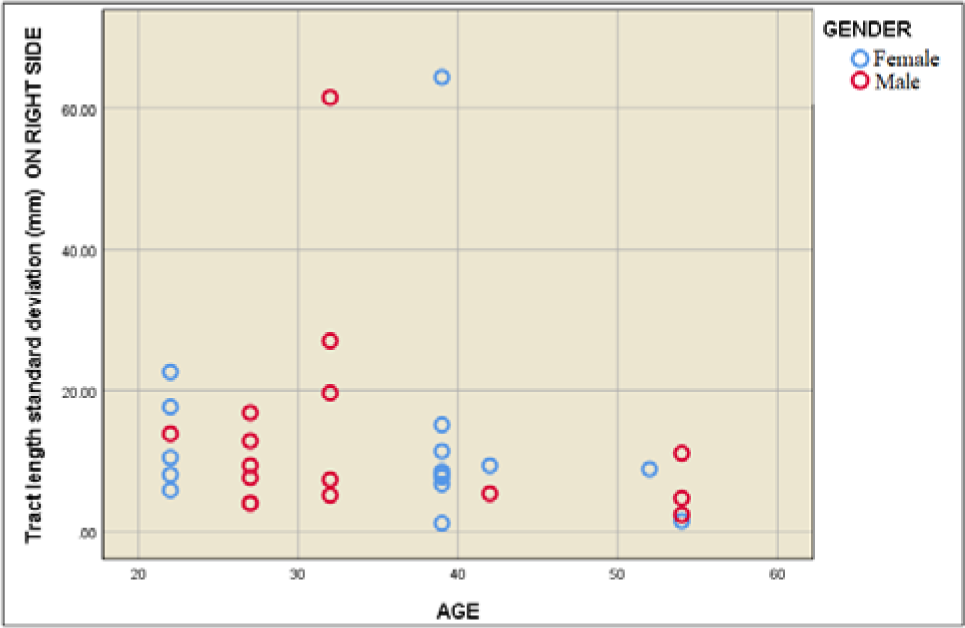
Graphical representation of tract length standard deviation (mm) on male and female subjects on right side.

**Table-32:** Comparison between Tract length standard deviation (mm) of both male and female unilateral hemispheric variances on left side *

| S.No. | Tract length standard deviation (mm) on left side in male subjects | Tract length standard deviation (mm) on left side in female subjects |
| --- | --- | --- |
| 1 | 6.11 | 37.53 |
| 2 | 5.35 | 23.21 |
| 3 | 4.76 | 2.49 |
| 4 | 3.12 | 8.26 |
| 5 | 46.99 | 8.19 |
| 6 | 13.59 | 3.01 |
| 7 | 6.38 | 6.44 |
| 8 | 39.99 | 4.91 |
| 9 | 5.29 | 4.29 |
| 10 | 2.95 | 57.09 |
| 11 | 6.30 | 5.90 |
| 12 | 4.32 | 5.83 |
| 13 | 5.23 | 77.31 |
| 14 | 79.59 | 2.81 |
| 15 | 5.92 | 4.88 |
| 16 | 7.77 | 3.01 |

\* The $p$ -value is .89432, result is statistically not significant at $p < .05$ .

**Graph-16:**
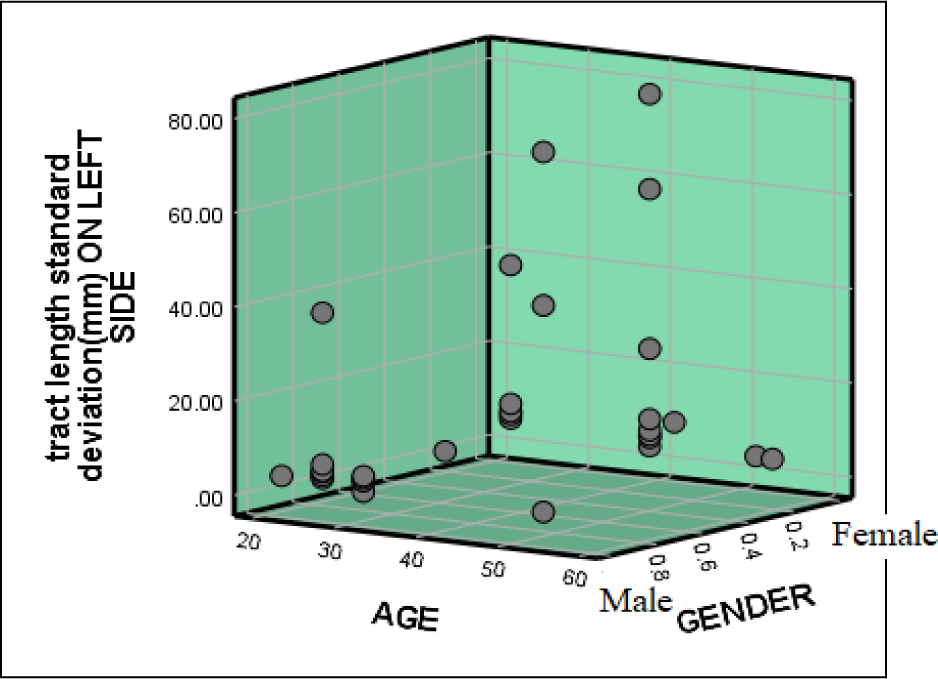
Graphical representation of tract length standard deviation (mm) from male and female subjects on left side.

## DISCUSSION

Saccades can be initiated by peripheral stimuli through uni-modal sensory interaction but are most times influenced by the interaction of multisensory signals where the potential target exudes signals from sensory modality interaction based on visual, auditory and even tactile inputs ^[14] [17] [23] [24] [26] [34] [35] [37] [43] [44]^. The influence of these associations on the production of saccades individually have been widely explored, predominantly establishing neural connectivity between auditory and visual pathways anatomically and functionally but such studies pertaining to olfactory-visual associations were yet to discover anatomical and functional links between both pathways ^[12] [16] [29] [30] [36] [38] [41]^. Auditory and visual sensory stimulations recorded by EEG in an experiment conducted by Kirchner, H. et al (2009) provided evidence that the FEF is not limited to eliciting saccades purely based on visual stimulations but found evidence that the FEF responded to auditory and visual stimulation within a similar scope suggesting that the FEF clearly processes multimodal signals ^[7] [11] [13] [15] [25] [31] [32] [33] [42]^. In light of this finding, we hypothesized and explored a possible structural connection between the FEF and the olfactory cortices.

Research studies previously conducted have proposed that visual stimuli influence olfactory attention through cross-modal interactions linked functionally through various pathways such as the dorsal and ventral visual streams ^[18] [22] [45] [50]^. A functional link between olfaction and the dorsal visual stream associated with motion perception and the ventral visual stream associated with object recognition have been found in previous studies but the neural structural connectivity and how the functional mechanisms interplay was still unexplored ^[9] [18] [19] [48]^. Based on studies carried out by Judauji, J. et al (2012) as well as another study by Morrot, G. et al (2001), both which investigated to what extent visual cues could affect olfactory processing through visual-olfactory associations, deduced that visual cues directly influences odor identification, perception and also attention ^[21] [51]^. Alternatively, studies conducted by Kuang, S. and Zhang, T. (2014) and another by Harvey, C. (2018) examined cross-modal integration in olfactory-visual coupling and found that olfactory cues influence gaze shifts and visual attention finding a functional pathway with which odor stimuli can initiate and guide eye movements ^[9] [49] [53]^. These studies made implications of functional associations between odor and visual pathways but the structural associations were still undiscovered ^[28] [39] [40]^.

Our data demonstrates a new finding that a structural connectivity exist between the ventral and dorsal Entorhinal cortex (Brodmann area 28 (V) & 34 (D), Piriform cortex (Brodmann area 27) and the Frontal eye field (Brodmann area 8), hence supporting previous documentation that suggested that olfactory cues could modulate visual attention, prompting saccades leading to spatial perception through neck and eye movements towards the stimuli ^[8] [9] [52]^. These new findings were discovered through non-invasive imaging methods where brain fibers were traced using diffusive tensor imaging from MRI imaging data collected from subjects both male and female and the results suggest that the finding of this study are mainly significant within the population and supported our initial conjecture. The data collected within the population generally showed consistent results in the parameters observed which were number of tracts, tract volume, tract length mean and tract length standard deviation in male and female subjects on both left and right brain hemispheres and for the most part, the difference in the data collected for males and females deemed insignificant. However, the slight variance was observed among some variables deviating from our subsidiary hypothesis where we suggested that general hemispheric similarities would be maintain when comparing between the parameters observed. It was observed among male subjects that on average, the right side had a greater number of tracts than the left side (graph-1) and it was also seen that female subjects had a greater number of tracts than male subjects on the left side (graph-4). These results have now unraveled a novel neuro-cortical pathway anatomically that fortifies studies that have previously indicated olfactory-visual coupling through cross-modal interaction between the senses ^[7] [8] [18] [19] [21] [22]^.

In converging the data of this study, the findings were deemed significant but maybe we were minimally limited in the sample size and diversity in randomness of collection of data within the population but since our findings were fairly novel, therefore in future research, a larger sample size could be explored with additional parameters and variables such as imaging of the neuropathology of the structural pathway we discovered. Also, since our topic of study is rather contemporary, literature to directly support the scope of our study was sparse and this report should then serve as a window to explore this topic further.

In light of these finding, additional functional and clinical correlations can be deduced as a potential biomarker to evaluate and assess certain neurodegenerative and psychiatric disorders where olfactory attention deficits are manifested as early onset of such disorders ^[2] [10]^. Most times olfaction is referred to as the vestigial sense and as one study showed, it is observed that olfactory function is given seldom evaluation in routine clinical examination hence minimal importance is placed on the fact that olfaction can provide conclusive assessments for cognitive function ^[2] [20] [27] [46] [47]^. A decline in olfactory function has been found to be an initial symptom of numerous neuro-degenerative such as Alzheimer’s disease, Parkinson’s disease, and others. Hence, early identification of olfactory functional deficit can serve as an initial symptom for differential diagnostic purposes with the ability to assess and evaluate strategies and therapies that can provide neuroprotective properties. This new structural connectivity found between the olfactory and visual pathway can be used in advancing neuroimaging technique for diagnostic purposes in evaluating cognitive decline and progression of disease ^[2] [10] [27]^.

In conclusion, this finding enhances our understanding of visual-olfactory associations and sensory coupling structurally in accordance with previous research that proposed its possible existence and corroborated its functional correlation. Olfactory dysfunction is known to be a preclinical sign of numerous neuro-degenerative diseases hence assessment of the structural integrity of this pathway could predict and deduce neurological decline providing diagnosis and prognosis of the patient’s condition. Confirmation of our finding can be further achieved through additional neuro-imaging analysis and this can ultimately direct additional appreciation to the usefulness of olfactory evaluation in the clinical setting.

Further research could focus on identifying and evaluating alterations in the structural integrity of the olfactory-visual pathway in patients that suffer from neurodegenerative diseases as it makes them susceptible to olfactory attention deficits and also the exact causes and pathological actions that lead to olfactory dysfunction in these diseases ^[2] [10]^. Methods such as functional MRI analysis could also assess the functionality of the odour-visual structural pathway that we found for further validation of our findings and then further implications such as establishments of evaluation methodologies revolving around olfactory attention could be implemented to serve as a clinical marker in diagnosing neurological disorders. Also, these finding should rivet more attention toward how critical olfactory function assessment should be in the clinical setting and in hopes that these findings are further supported since olfactory function evaluation could possibly predict neuro-aging deficit predisposition and assess cognitive decline.

## MATERIALS AND METHODS

### Datasets Acquisition

The study used an open access, ultra-high b-value, various diffusion sensitizing directions (64,128 and 256) diffusion imaging datasets with the in-plane resolution and slice thickness of 1.5 mm. The datasets are originally developed and reposted by Massachusetts General Hospital – US Consortium Human Connectome Project (MGH-USC HCP).

The present study involves Thirty-two healthy adult datasets (16 males and 16 Females, between the age 20–59 years old; mean age = 30.4). Given de-identification considerations, age information is provided in 5-year age bins (Table-33). All participants gave written informed consent, and the experiments were carried out with approval from the Institutional Review Board of Partners Healthcare of MGH-USC HCP project [80]. The demographic details of the participants with gender and age are available in the data sharing repository.

**Table 33:** Age information of male and female subjects

| Subject No. | Age in years | Number of Male subjects | Number of Female subjects |
| --- | --- | --- | --- |
| 1 | 20-24 | 1 | 5 |
| 2 | 25-29 | 6 | 8 |
| 3 | 30-34 | 5 | 0 |
| 4 | 35-39 | 3 | 1 |
| 5 | 40-44 | 2 | 1 |
| 6 | 45-59 | 2 | 1 |
|  | TOTAL | 16 | 16 |

### Data Pre-processing and Fibre tractography

The MGH-USC HCP team has completed their basic imaging data pre-processing, with software tools in Free surfer and FSL, which includes i. Gradient nonlinearity correction, ii. Motion correction, iii. Eddy current correction, iv. b-vectors. Fiber tractography is a very elegant method, which can used to delineate individual fiber tracts from diffusion images. The main process of study uses the “DSI-Studio” software tools for i. Complete pre-processing, ii. Fiber tracking and iii. Analysis.

## ACKNOWLEDGEMENTS

We thank our colleagues from “Team NeurON” for their insights and assistance in compiling the manuscript of our research paper.

